# North American Birds Require Mitigation and Adaptation to Reduce Vulnerability to Climate Change

**DOI:** 10.1101/798652

**Authors:** Brooke L. Bateman, Chad Wilsey, Lotem Taylor, Joanna Wu, Geoffrey S. LeBaron, Gary Langham

## Abstract

Biodiversity is being lost at an alarming rate across the globe, with extinction rates up to a hundred times greater than historical norms. Climate change will only exacerbate this crisis. The rapid pace of projected climate change is set to push birds to seek new locations, drastically reshuffling the avian communities of North America. In an emerging climate crisis, effective conservation requires both adaptation and mitigation to improve the resilience of species. However, the pledged reductions in greenhouse gas emissions outlined in the Paris Agreement framework would still lead to a 3.2°C or greater increase in global mean temperature by the end of this century. In this study, we use big data analytics to develop species distribution models and assess the vulnerability of 604 North American birds to multiple climate change scenarios. We assess how climate change mitigation can affect the number of species vulnerable to climate change, as well as the species and locations at risk if emissions continue unchecked. Our results indicate that over two-thirds of North American birds are moderately or highly vulnerable to climate change under a 3.0°C global warming scenario. Of these climate-vulnerable species, 76% would have reduced vulnerability and 38% of those would be considered non-vulnerable if warming were stabilized at 1.5°C. Thus, the current pledge in greenhouse gas reductions set by the Paris Agreement is inadequate to reduce vulnerability to birds. Additionally, if climate change proceeds on its current trajectory, arctic birds, waterbirds and boreal and western forest birds will be highly vulnerable to climate change; groups that are currently not considered of high conservation concern. Thus, there is an urgent need for both aggressive policies to mitigate emissions and focused conservation adaptation actions to protect birds and the places they need in a changing climate.

## INTRODUCTION

Biodiversity is being lost globally at an alarming rate, with extinction rates up to a hundred times greater than historically seen [1]. Within North America, 37% of bird species are under high risk of extinction [2]. Climate change exacerbates the global biodiversity crisis [3], with 24–50% of bird species vulnerable to climate change alone [4], and 11–15% both vulnerable to climate change and already threatened with extinction per the IUCN Red List. The threat of climate change affects species differently, as each has a unique set of physiological tolerances, competitive interactions, genetic variation, and habitat requirements. These differences translate into range limits for species and across species into patterns of biodiversity. Across landscapes, climate and weather have been shown to limit range boundaries [5–7]; regulate abundance [8], affect spatial patterning [9] and species richness of birds [10]; and directly influence habitat quality and type [11]. Given that climate has a strong influence on the spatial patterns of bird occurrence and abundance, changes in climate will disrupt both species and communities.

Given enough time, species can, and historically are known to, respond to climate change by shifting their ranges and tracking suitable environmental conditions as they move across the landscape [12,13]. Indeed, birds already respond to contemporary climate change with range shifts [14–19]. However, on our current trajectory, contemporary climate change is anticipated to be 20 times faster in the next century than at any period over the last two million years [20], and even common and wide-spread species are anticipated to experience significant range contractions [18]. This rapid pace of climate change will lead to more range shifts [21], pushing birds to seek new locations at a greater velocity [22], and drastically reshuffling the avian communities of North America [16,23]. Effective climate change conservation requires an understanding of species’ vulnerability to climate change. Vulnerability is based on species’ specific exposure, sensitivity, and adaptive capacity [24–27]; e.g. the local pressures to be experienced from a changing climate, the characteristics that result in sensitivity to change, and how well a species may be able to cope with change, respectively. Here, we assess climate change vulnerability of North American bird species. Armed with this information, we can better gauge where to focus conservation efforts for birds in a changing climate.

Furthermore, we utilize our understanding of vulnerability to inform climate change mitigation and adaptation to reduce vulnerability and improve species’ ability to cope with climate change [24,28,29]. Identifying appropriate mitigation, policy actions set to reduce the magnitude of climate change as much as possible, and adaptation, conservation actions to help cope with the consequences of inevitable climate change [30], strategies are thus imperative.

To understand how climate mitigation could potentially alter species vulnerability to climate change, we assess vulnerability of birds in North America in the context of accepted future climate trajectories. The Intergovernmental Panel on Climate Change (IPCC) recommends a limit in global mean temperature to less than 2.0°C above pre-industrial levels, and preferably to 1.5°C if possible [31–34]. Currently, we are on track to surpass this limit: ∼1.0°C (0.8-1.2°C) of warming has already occurred [32] and 3.0°C is expected by 2100 under a conservative business-as-usual trajectory [35]. Indeed, there is high confidence that we will reach 1.5°C between 2030 and 2052 [32] regardless of emissions reductions strategies. The currently pledged reductions in greenhouse gas emissions (i.e., Nationally Determined Contributions) outlined in the Paris Agreement framework would lead to at least a 3.2°C increase in global mean temperature [35]. Here we assess the risk of climate change vulnerability to birds in North America for three policy-relevant scenarios: 1.5°C, 2.0°C, and 3.0°C increases in global mean temperature. These scenarios reflect current and potential climate change emissions reduction targets and provide relevant content for national policies that would address mitigation of the threat of climate change.

To address optimal adaptation strategies and identify where we can focus conservation actions to help buffer species from rapid change, we next determine which bird species and which locations in North American are at risk under possible climate change scenarios. High-resolution, continental scale projections of species range shifts from species distribution models (SDMs) are able to provide information at the spatial scale relevant to on-the-ground actions. SDMs can give us a snapshot of potential futures to help identify which scenarios will be problematic, and which regions and species might be at risk. Birds serve as an ideal study group to forecast potential realities in a changing climate using SDMs [36]. First, they are ubiquitous taxa, and the millions of geo-located observations covering much of North America are now available in global databases. These avian data sets come from the growth of community science programs such as eBird; large online databases such as the Global Biodiversity Information Facility (GBIF), Avian Knowledge Network (AKN), and Biodiversity Information Serving Our Nation (BISON); long-term surveys, such as the North American Breeding Bird Survey (BBS); and a multitude of scientific studies by individual researchers. Second, birds serve as a conservative baseline for other taxa facing climate change. Given their higher dispersal rates and migration capacity, it is likely that birds will fare better than less vagile species [37,38]. Paired with the continental-scale availability of high-resolution environmental data, remote sensing products, and future projections of climate and vegetation change, and with a well-developed field of SDM research [39], these data sets allow us to provide projections of future possibilities for bird species that are relevant for adaptation planning [39,40].

In this work, we compile more than 140 million observations to develop SDMs and an assessment of climate change vulnerability for 604 North American bird species at a continental scale in both breeding and non-breeding seasons and under multiple global climate change scenarios. We aim to identify how different climate change targets translate to potential changes in bird communities across North America in order to aid in both mitigation policy and conservation adaptation decisions to ameliorate the impacts of global climate change on birds.

## MATERIALS AND METHODS

We assess species’ vulnerability to climate change as a function of a species’ climate change exposure, sensitivity, and adaptive capacity [27]. We do so for 604 species across Canada, the United States, and Mexico using a combination of species distribution models (SDMs) and trait-based information [41]. In doing so, we adapt the methods from Wilsey et al. (2019) to develop mitigation policy and conservation adaptation relevant results. Our modeling effort includes data extraction and filtering, model building and evaluation, and model projection under climate change [25].

### Response Variable: Bird Data

We compiled more than 140 million bird occurrence records from 70+ datasets across Canada, the United States, and Mexico for 604 species in the breeding and non-breeding seasons. We cleaned all datasets to remove occurrence records that were incomplete (i.e., missing geographic coordinates), extraneous (i.e., coordinates outside of the survey area of the data source), had inexact spatial information (i.e., traveling surveys longer than 1 km or area surveys greater than 100 ha), were of long durations (>180 min), or were greater than 1 km in extent. For data from GBIF, we only downloaded data not included in other datasets (e.g. eBird). To account for variation in species identification across multiple datasets with varying expertise on bird taxa, we took several additional approaches to clean our data and remove unreasonable records. This included a vagrant rule to remove records for rarely occurring species and a minimum of two experts to review each species/season occurrence dataset. First, we established a minimum ‘vagrant rule’, requiring a species to occur in a Bird Conservation Region (BCRs; NABCI, http://nabci-us.org/resources/bird-conservation-regions-map/) in at least two separate years. For some species, we increased this vagrant rule to drop BCRs that contained vagrant records in more than two years. Next, we visually inspected each species occurrence dataset in both the breeding and nonbreeding season and dropped erroneous records from BCRs outside of the current range of the species (based on bird range maps from the Birds of North America [42]) if increasing the vagrant rule did not sufficiently address this issue. Finally, we enlisted a minimum of two experts to review each species/ season occurrence dataset and identify erroneous records. Others have identified that expert opinion improves species distribution models [43]. In some instances, experts identified additional records to remove that were not contributors to the breeding populations (e.g. immatures, ‘lame’ or non-migrating individuals, or feral stock).

We included bird species where their breeding or non-breeding ranges occurred primarily within the United States and Canada [36], and included all available bird occurrence data across North America for these 604 species in order to sample from the full range of environmental conditions possible for each species [39,44]. These data included a combination of both structured and unstructured avian surveys, so we took action to reduce sampling biases that are inherent in non-systematically designed surveys. To address bias and avoid model over-fitting [45,46], we applied both a geographic filtering approach and a target-group background approach [47], weighting our background sampling by the number of occurrence points within each grid cell. To ensure that the geographic extent of our data was appropriate, we defined the extent from which we sampled both the occurrence and background points using a movement-hypothesis approach to improve model performance and generalizability [45]. This approach accounts for regions that a species could have experienced historically through movement [44] and are potentially suitable and accessible, but do not include areas beyond this [39]. We extracted background points with an approximate ratio of 5:1 background points per occurrence points, but restricted the number of points to be between 10,000 and 100,000, with a minimum density of 0.0001 points/ km2 in each Bird Conservation Region (BCR). For some species, we manually added or dropped BCRs from background sampling when selecting adjacent BCRs was inconsistent with a species’ natural history. See S1 Bird Data Sources for a full list of datasets sources.

To distinguish breeding and non-breeding occurrence records, we first assigned each species a resident status based on published range maps [42], life history [42] and expert opinion. Species were classified as breeding-only (non-breeding range outside of North America), two-season (separate breeding and non-breeding ranges within North America), or permanentresident (non-migratory). We consulted with experts to define default breeding (June-July) and non-breeding (January-February) seasonal date ranges that worked well for most species, and custom seasonal date ranges for species with different migration timing (breeding season of June only for plovers, sandpipers, and hummingbirds; wintering season of December only for hummingbirds and swallows, December-January for grassland birds). We assigned each species to a habitat affinity group based on NABCI’s 2009 State of the Birds report (www.stateofthebirds.org). These included: arctic, aridlands, boreal forests, coastal, eastern forests, generalists, grasslands, marshlands, subtropical forests, urban/suburban, waterbirds, and western forests (range in the number of species per group = 9-89). For species encompassing more than one group, we selected the group that best matched their habitat needs. In a few instances, we grouped species in different habitats for breeding and non-breeding seasons based on their ecology (see S5).

### Predictor Variables: Climate and Environmental Data

#### Climate Data

We used gridded current and modeled future climate developed by AdaptWest as covariates in the SDMs [48]. Climate grids consisted of 23 million grid cells covering North America at a 1 km resolution. For the present, we used statistically downscaled climate normals for the time period of 1981-2010 derived from the Climatic Research Unit Time series 3.22 dataset [48]. For future projections, we used Coupled Model Intercomparison Project phase 5 (CMIP5) projections from three individual General Circulation Models (GCMs) and an ensemble of 15 GCMs. The three individual GCMs capture a range of intermediate (CCSM4), warm-wet (GFDL-CM3), and cold-dry (INM-CM4) future conditions for the continent [48]. We included two greenhouse gas representation concentration pathways (RCPs; RCP 4.5 and RCP 8.5) for two future time periods (2050s and 2080s). To provide climate scenarios relevant to policy, we associated a 1.5°C global mean temperature rise with RCP 4.5 for the 2050s, 2.0°C with the RCP 8.5 for the 2050s, and 3.0°C with the RCP 8.5 2080s. Multi-year climate averages have been shown to be a sufficient temporal scale to capture species ranges and projections for species distribution models [49].

The final climate covariates, after reduction of collinearity, included nine bioclimatic variables; this included mean temperature of the warmest month, chilling degree days (degree-days below 0°C), and summer heat moisture index for the breeding season. For the non-breeding season, we included mean temperature of the warmest month, and growing degree days (degree-days above 5°C.) In addition to these season-specific climate variables, we included climatic moisture deficit, number of frost-free days, mean annual precipitation, and precipitation as snow for both breeding and non-breeding seasons. For more details on climate data and selection, see Wilsey et al. [25].

#### Environmental Data

We included key environmental variables to capture habitat-specific information based on ecological knowledge of North American bird species. We included proximal predictor variables beyond climate for all species, based on habitat group. For all species, we included vegetation type [50], terrain ruggedness [51], and anthropogenic land cover [52]. For specific habitat groups, we used ecologically relevant variables such as surface water for waterbirds and marshbirds [53], wetland type for waterbirds and marshbirds [54], distance to wetlands for waterbirds, distance to coast (excluding inland water bodies) for coastal birds, distance to shore (including inland water bodies) for marshbirds and waterbirds [55], and a human influence index for urban/suburban birds [56].

Most habitat variables were only available for the present, except vegetation type. Future vegetation projections represented a consensus across three GCMs (CGCM3, HadCM3, GFDL CM2.1) and two Special Report on Emissions Scenarios (SRES) emissions scenarios (A2 and B1 or B2) from CMIP3, as outlined in [50]. CMIP3 and CMIP5 projections are more similar than distinct [57] and for these purposes represent the best available vegetation projections on the continental scale [25]. We used the mid-century vegetation projections to align with the 1.5°C and 2.0°C, and late-century projections for the 3.0°C global mean temperature scenarios. For all other habitat variables, current conditions were included in future projections as the best available information.

### Data Processing

We aligned covariate data to a common grid covering our study area at a 1-km resolution in an Albers equal area conic projection using utilities from GDAL/OGR [58]. We resampled data with different methods to account for data type and original resolution, using bilinear interpolation for high-resolution (∼1 km) continuous data (climate variables, human influence index), median (terrain ruggedness) or max (surface water) for higher-resolution (< 1 km) continuous data, nearest neighbor for high-resolution categorical data (vegetation type), and mode for higher-resolution categorical data (anthropogenic landcover, wetland type). To expedite data queries connecting environmental data with corresponding bird occurrence data, we built a PostgreSQL relational database organized around a common and unique grid ID field. See Wilsey et al. (2019) for comprehensive details on data processing. Our 1-km format provides high-resolution outputs represents a spatial scale to which birds ranges are regulated by climate and broad-scale habitat features (citation). [59,60]

### Species Distribution Modeling

#### Model Building

We approached our modeling effort by integrating the latest modeling approaches with the natural history of each species, with the goal of producing the best model for each species instead of a one size fits all approach [61]. Our modeling effort includes data extraction and filtering, model building and evaluation, and model projection under climate change [25]. We modeled each species within a habitat group context, using ecologically relevant variables for each group. We built both breeding season and non-breeding season models for two-season and resident species to capture the seasonal differences in bird species ranges. We also included a multi-step expert review process where we had an expert assess the bird occurrence data, modeled current range, and projected future range for each species and season [43].

#### Model Parameters and Performance

We used an internal repeated geographical sub-sample of training data to generate a robust assessment of model performance for model selection. We performed internal cross-validation within our model building based on a 10-fold 75:25 percent training-to-testing data partition using a masked geographically structured approach [62]. To inform this, we used semivariograms to identify the spatial scale that variability in our environmental data leveled off at (200 km) [25]. We then applied a 200-km grid over our study area and randomly sampled our occurrence data into 10 representative training and testing datasets [63]. Each of these datasets were held constant for all models for a species to ensure comparability of performance measures, and to maintain constant prevalence between the training and testing datasets [64].

We built models with two algorithms, boosted regression trees (BRTs) and maximum entropy (Maxent, version 3.3.3k), implemented in the dismo package in R. Both approaches are well regarded in the literature for accuracy and their ability to model non-linear species-habitat relationships [62,65]. We first built BRTs to evaluate an appropriate filtering resolution (i.e. 1, 10, or 50 km). Model performance was evaluated using area under the receiving operating characteristic curve (AUC) for predictions to the spatially stratified training datasets. We compared filtering scenarios using median AUC and the top-performing filtering approach to build Maxent models. We built Maxent models that varied the regularization parameter from 0.5 to 4.0 and compared model performance with median AUC as above. As our goal was projection under future climate change scenarios, we aimed to minimize generalizability and reduce overfitting [46] by selecting the top-performing model based on median AUC across all BRT and Maxent model runs. We then used the top model for model prediction.

#### Model Evaluation

We evaluated multiple thresholds for each species and season. We calculated our threshold choices using the SDMTools package in R [66]; these included mean occurrence prediction (mo; mean suitability prediction for the occurrence records), maximum sensitivity specificity (tss; maximized sensitivity + specificity), 10% omission (om_10; excludes 10% of occurrence records), sensitivity specificity (eq; sensitivity is equal to specificity), maximum Kappa (mk; maximum Kappa statistic), minimum occurrence prediction, (min_pred; minimum suitability prediction across all occurrence records), and a derived custom threshold to fall between the minimum occurrence prediction and 10% omission thresholds ((om_10+ min_pred)/3). For each threshold applied, we calculated the true positive (commission error) and true negative (omission error) rate for how each model classified presence and background points. After visual inspection of each model output, a final threshold was included that aligned with expert opinion and minimized a decrease in model performance. To ensure models approximated ecological reality, we then evaluated each species and season output through an expert review process. This included a first pass comparison of models with range maps [42], followed by a rigorous external review by at least two experts for each bird species. Where species models deviated from known ecology (e.g., errors in occurrence records in location or representation of predictor variables, modeled range errors, over- and under-prediction due to unrealistic dispersal projections), efforts were taken to remodel. Steps included for re-modeling included dropping erroneous occurrence records, adding or dropping BCRs from the species background, or changing the selected threshold.

#### Model Projection

We generated high-resolution (1 km) predictive occurrence maps for each season and scenario (1.5°C, 2.0 °C and 3.0 °C). To provide a range of potential futures, for each scenario we projected onto an ensemble as well as three individual GCMs that included a range of scenarios of future climate change including warm-wet, and cold-dry (as detailed above). To account for uncertainty from novel conditions, we limited extrapolation by only projecting onto vegetation and land-use classes that were included in the model training datasets for each species. We also use the ‘clamping’ feature in Maxent and Boosted Regression Trees to limit projections to novel climates. For resident and breeding season models, we estimated dispersal limitations based on mean natal dispersal [67] and generation time [68] for each species [67]. We did not include dispersal limitations for non-breeding season models, as there is limited information available on non-breeding movement and site fidelity for the majority of bird species. To address model commission error and limit biological extrapolation, we applied an expert-opinion based approach that identified BCRs with over-prediction [43]. We masked current and future projections from BCRs with commission error from both current and future projections if it was identified that the BCRs were geographically distinct and/or biological limitations makes it unlikely for the species to disperse there.

### Vulnerability Assessment

To evaluate mitigation strategies, we assessed species vulnerability to climate change under multiple climate change scenarios. Vulnerability to climate change is a combination of each species’ exposure, sensitivity, and adaptive capacity [24–27]. Here, we define exposure for each species as model-based climate change projections of species ranges for multiple climate chance scenarios, sensitivity as how much of a species’ range is lost under future climate change in each scenario, and adaptive capacity as the ability of a species to buffer from climate change through shifting or gaining range in each scenario. Our possible exposure scenarios reflect relevant climate change mitigation policy target scenarios, with a 1.5°C global mean temperature rise as potential climate stabilization scenario, 2.0° C as in intermediate scenario based on IPCC recommendations, and a 3.0°C scenario as our current business-as-usual estimate. In each of our future scenarios, we compared current and future range maps to classify areas projected to be lost, gained, or maintained for each species and season. Areas that are currently suitable but become unsuitable (i.e., drop below the threshold) are considered loss, areas that are currently unsuitable but become newly suitable (i.e., pass above the threshold) are considered gain, and areas that are suitable in both periods (i.e., remain above the threshold) are considered stable. We assessed vulnerability based on projected range loss and potential gain [25,36,69] per Wilsey et al. (2019). We use the proportion of a range loss to define climate sensitivity, and the ratio of range gain to range loss to define adaptive capacity. We classified climate sensitivity by binning proportion range loss into four equal intervals (0-25%, 25-50%, 50-75%, 75-100%) and assigning a value of 0-3 from low to high across bins. We classified adaptive capacity by binning the ratio of range gain to range loss into four classes (>2:1, 1-2:1, 0.5-1:1, and 0-0.5:1) and assigning a value of 0-3 from low to high across the bins. We then summed the climate sensitivity and adaptive capacity scores to get the final vulnerability score for each species and season, classifying them as neutral (sensitivity + adaptive capacity = 0), low vulnerability (sensitivity + adaptive capacity = 1 or 2), moderate vulnerability (sensitivity + adaptive capacity = 3 or 4), and high vulnerability (sensitivity + adaptive capacity = 5 or 6). We classified the final vulnerability class for each species and season, classifying them as neutral, low vulnerability, moderate vulnerability, and high vulnerability. With this approach, species that experience greater range loss without being able to make up for it in range gain have higher vulnerability. Here, we consider species in the moderate and high vulnerability classes to be vulnerable to climate change. To calculate confidence in our vulnerability scores, we assessed the number of times the vulnerability score agreed across the ensemble model and the three individuals GCMs for each of the future scenarios.

We also assess the number of species that we classify as climate vulnerable compared to the species currently listed on the Partners in Flight (PIF) Watch lists [70], a list of species currently considered of highest conservation concern across North America. PIF Watch list data was available for 495 of our 604 species.

### Spatial Assessment

To transfer the species distribution projections into tools to inform climate change adaptation, we mapped changes in bird community composition across North America [40,71]. We mapped areas of aggregated species loss, gain, and net change in community composition for each of the habitat-associated species groups and for all species. Climate change adaptation strategies can be tailored for areas with high projected loss (local extirpation), gain (colonization), or net change [72].

## RESULTS

Nearly two-thirds, 64% (389/604), of species were climate vulnerable in at least one season and scenario (S4 Table). Some habitat groups were highly vulnerable across the majority of species (Fig 1), including 100% of arctic birds (16/16), followed by 98% of boreal forest birds (47/48), 86% of western forest birds (63/73), and 78% of waterbirds (66/85). Habitat groups with intermediate vulnerability included subtropical forests (71%, 25/35), grasslands (68%, 27/38) eastern forests (59%, 41/69), and coastal (52%, 31/60). Habitat groups with lower vulnerability included urban/suburban (50%, 3/8), aridlands (45%, 31/69), marshlands (41%, 25/61), and generalists (31%, 15/48), although even in these groups more than a quarter of the species were considered climate vulnerable.

**Fig. 1.**
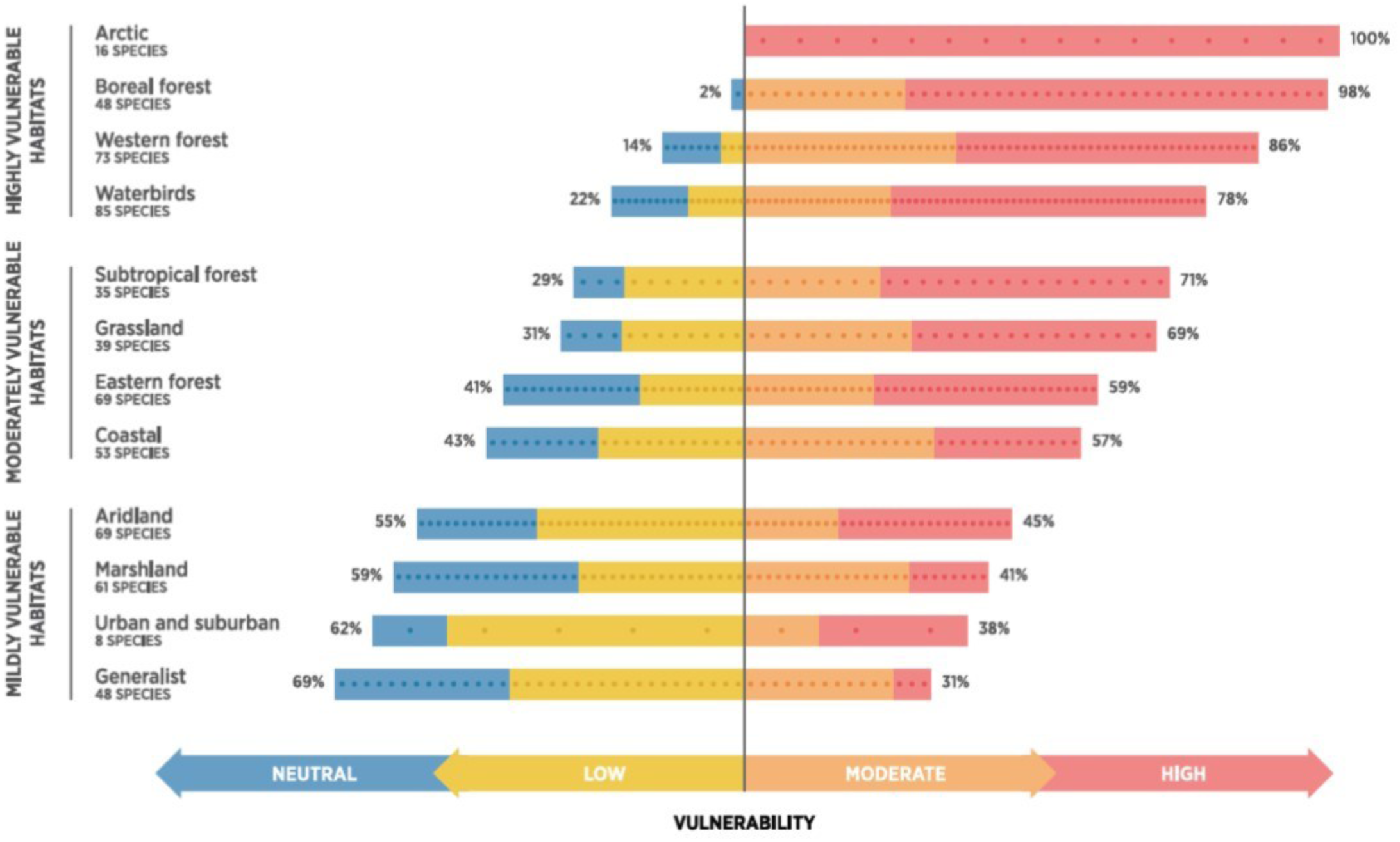
Percentage of species vulnerable by bird habitat groupings across all scenarios and seasons. Purple and green indicate percentage of species vulnerable (high and moderate) on a positive axis, and blue and yellow indicate non-vulnerable (low and neutral) on a negative axis.

Vulnerability declined in scenarios with greater mitigation to reduce warming (Fig 2). Nearly two-thirds (63%, 383/604) of all species across both seasons were classified as vulnerable at 3.0 °C, compared to 54% at 2.0°C (N = 327), and 47% at 1.5°C (N = 286) global mean temperature rise (Fig 2). In fact, across seasons for species considered vulnerable (high and moderate vulnerability) at 3.0°C, 76% (290 unique species of 383, 356 combinations of breeding and non-breeding) drop at least one climate vulnerability category lower, and 38% (146 unique species of 383, 159 combinations of breeding and non-breeding) are no longer vulnerable under 1.5°C. One species, the Golden-cheeked Warbler (Setophaga chrysoparia), could drop vulnerability by up to three categories, from high vulnerability at 3.0° C, to neutral vulnerability at 1.5°C in the breeding season.

**Fig. 2.**
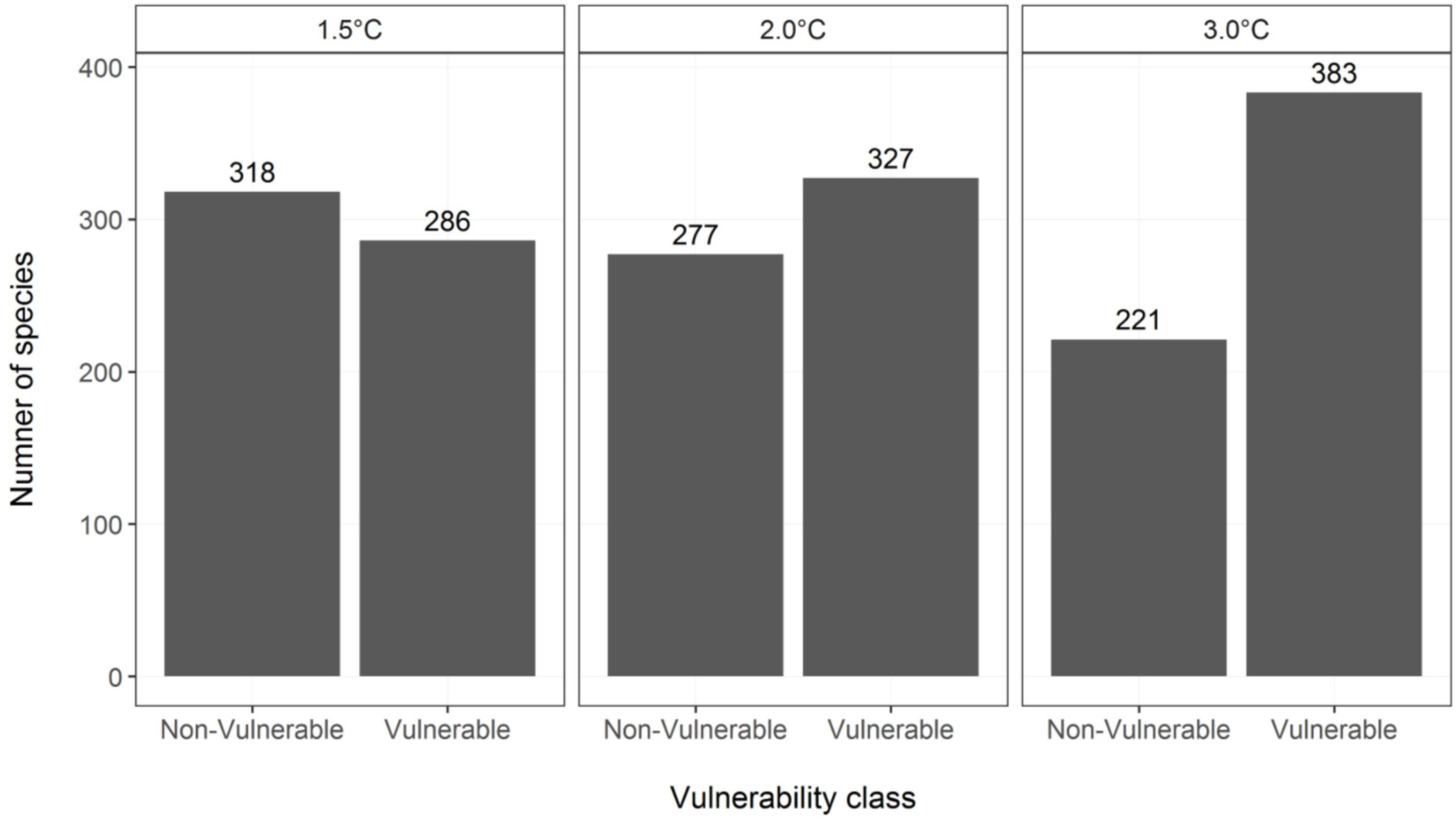
Number of species classified as vulnerable or non-vulnerable under 1.5°C, 2.0°C and 3.0°C global warming scenarios. Vulnerable species are within the moderate or high vulnerability classes, and non-vulnerable species are within the neutral or low vulnerability classes.

Vulnerability was considerably higher in the breeding season compared to the non-breeding season. In the breeding season, 58% (345/597) of species are vulnerable at the 3.0°C scenario. However, stabilizing global warming at 1.5°C global mean temperature rise would bring this down to 43% (254/597) of breeding species classified as vulnerable (Fig 3, S4 Species Vulnerability). In the breeding season, the most vulnerable groups at 3.0°C were arctic (100%) boreal forests (98%), western forests (78%), and waterbirds (78%) (S3.1 Table). In these four groups, at least 62% of species were still vulnerable at 1.5° C in the breeding season. Stabilizing mean global temperatures at 1.5°C as compared to 3.0°C had the greatest reduction in the number of species vulnerable in the breeding season, the most for the urban/suburban group (38% reduction), followed by the boreal forests (23% reduction), grasslands (23% reduction), eastern forests (20% reduction), and generalists (17% reduction) groups. We saw moderate reductions in the vulnerability of western forests, subtropical forests, marshlands, aridlands, waterbirds, and coastal species in the breeding season, with additional climate change adaptation actions required for arctic species, which were 100% vulnerable regardless of climate change scenario (See S3 Supplementary Figures and Results, S3.1-12 Figs).

**Fig. 3.**
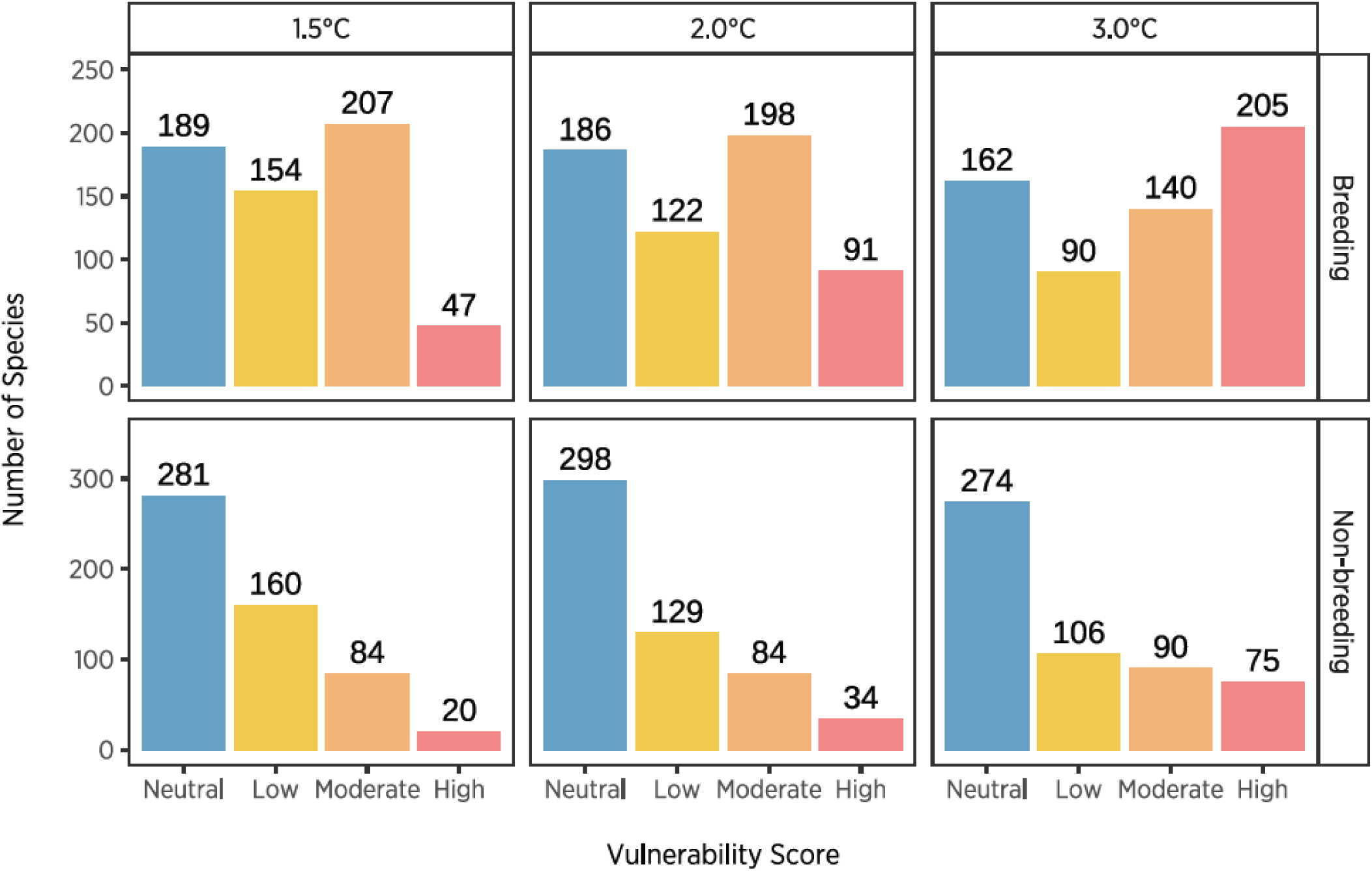
Vulnerability classification in the breeding and non-breeding seasons under 1.5°C, 2.0°C and 3.0°C global warming scenarios. Vulnerable species are within the moderate or high vulnerability classes, and non-vulnerable species are within the neutral or low vulnerability classes.

Vulnerability was considerably lower in the non-breeding compared to the breeding season, with 30% (165/546) of species in the non-breeding season vulnerable at the 3.0°C scenario. This would be reduced to just 19% (104/546) of species classified as vulnerable in a 1.5°C scenario in the non-breeding season. In the non-breeding season, the most vulnerable groups include subtropical forests (59%), boreal forests (58%), western forests (51%), and arctic (47%) species (S3.1 Table), although vulnerability was considerably lower across all groups compared to the breeding season. All groups had reduced vulnerability in the 1.5°C scenario, with the greatest effects seen in boreal forests (24% reduction) and grasslands (20% reduction) in the non-breeding season (S3.1 Table). Across all species and scenarios, we found 93% agreement (48% agreement across all scenarios, 45% across three scenarios) on vulnerability scoring across our GCM ensemble and three individual GCM scenarios. Vulnerability scores for all species and model agreement across our GCM ensemble and three individual GCM scenarios can be found in S4 Species Vulnerability.

Areas of overlapping range loss across species varied by both scenario (1.5 vs 3.0°C) and season (breeding vs. nonbreeding; Fig 4). In the breeding season, bird communities lost up to 106 species locally with a 3.0°C scenario, with a pronounced loss of species in the northern and eastern temperate forests, the Boreal Shield of Canada, and the northern parts of the Midwest and Northeast in the United States. Although less concentrated, losses were also high in the Pacific Northwest, Rocky Mountains, and Alaska. Losses were less dramatic under a 1.5°C scenario, with up to 79 species lost locally, and concentrated mostly within the Boreal Plain region of Canada. In the non-breeding season, species losses were lower, with up to 70 species lost from communities locally at 1.5°C scenario and 90 at 3.0°C, spread out across the United States and Mexico. High loss is seen in the Yucatán Peninsula and eastern Mexico at 1.5° C, with patterns shifting to more pronounced loss in the western US and the Canadian boreal under the increased 3.0°C warming scenarios (Fig 4).

**Fig. 4.**
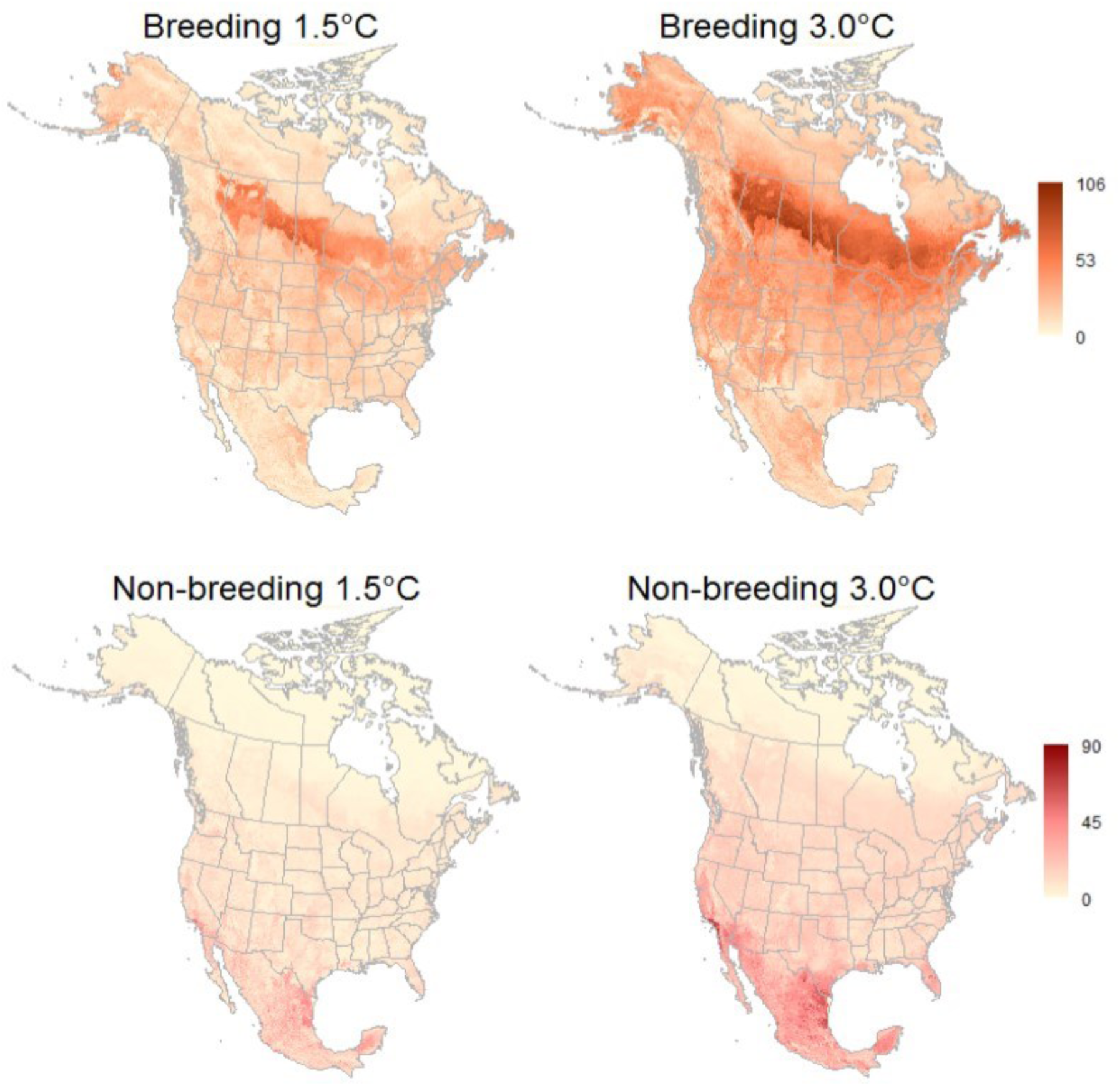
Tallies of overlapping species with projected range loss across North America in the breeding and non-breeding seasons under 1.5°C (left) and 3.0°C (right) warming. The number of species with overlapping range loss in a single grid cell in the breeding season ranged from 0-106, and from 0-90 in the non-breeding under 3.0°C warming. Under 1.5°C warming, counts of projected range loss in the breeding range from 0-79, and from 0-70 in non-breeding.

Communities gained up to 115 potential colonizers per grid cell in the breeding season and 105 in the non-breeding season under the 3.0°C scenario (Fig 5). In the breeding season, peak gains were concentrated in the taiga and tundra ecoregions of Canada, and to a lesser extent in the northern forests of the United States and Canada and along mountain ranges like the Rockies and Sierra Madre (Fig 5). In the non-breeding season, peak gains were concentrated in western Alaska, Newfoundland, Nova Scotia, the Great Lakes, the central southern US, and the Sierra Madre in Mexico. In both the breeding and non-breeding seasons, species range gains are more dilute at the 1.5° C scenario with no peak concentration areas identified.

**Fig. 5.**
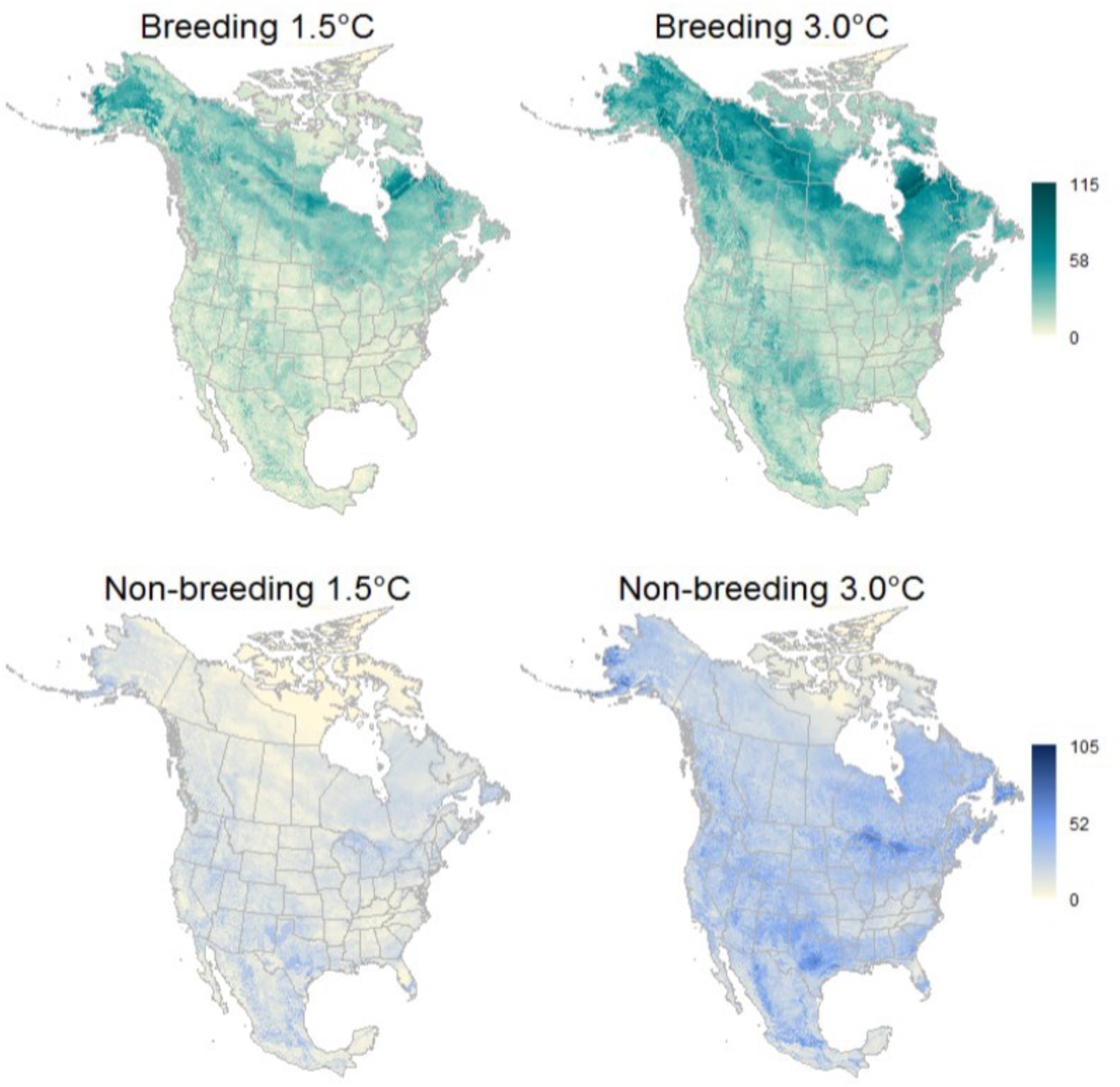
Tallies of overlapping species with projected range gain across North America in the breeding and non-breeding seasons under 1.5°C (left) and 3.0°C (right) warming. The number of species with overlapping range gain in a single grid cell in the breeding season ranged from 0-115, and from 0-105 in the non-breeding under 3.0°C warming. Under 1.5°C warming, counts of projected range loss in the breeding season ranged from 0-93, and from 0-66 in non-breeding.

We projected a net decline in species richness across most of North America in both scenarios for the breeding season, except in the tundra and taiga ecoregions, the Southern Great Plains in Texas and New Mexico, and parts of the Rocky and Sierra Madre mountains. Net losses were more widespread in the US and the boreal region of Canada under the 3.0°C scenario, but is still prevalent at 1.5°C (Fig 6). In contrast, the non-breeding season was projected to experience limited net loss across all of North America in either scenario, with net loss restricted to Mexico, Florida, and southern California (Fig 6). Projected gains in the non-breeding season were greatest under 3.0°C in western Alaska, Newfoundland, Nova Scotia, the Great Lakes, and central Texas.

**Fig. 6.**
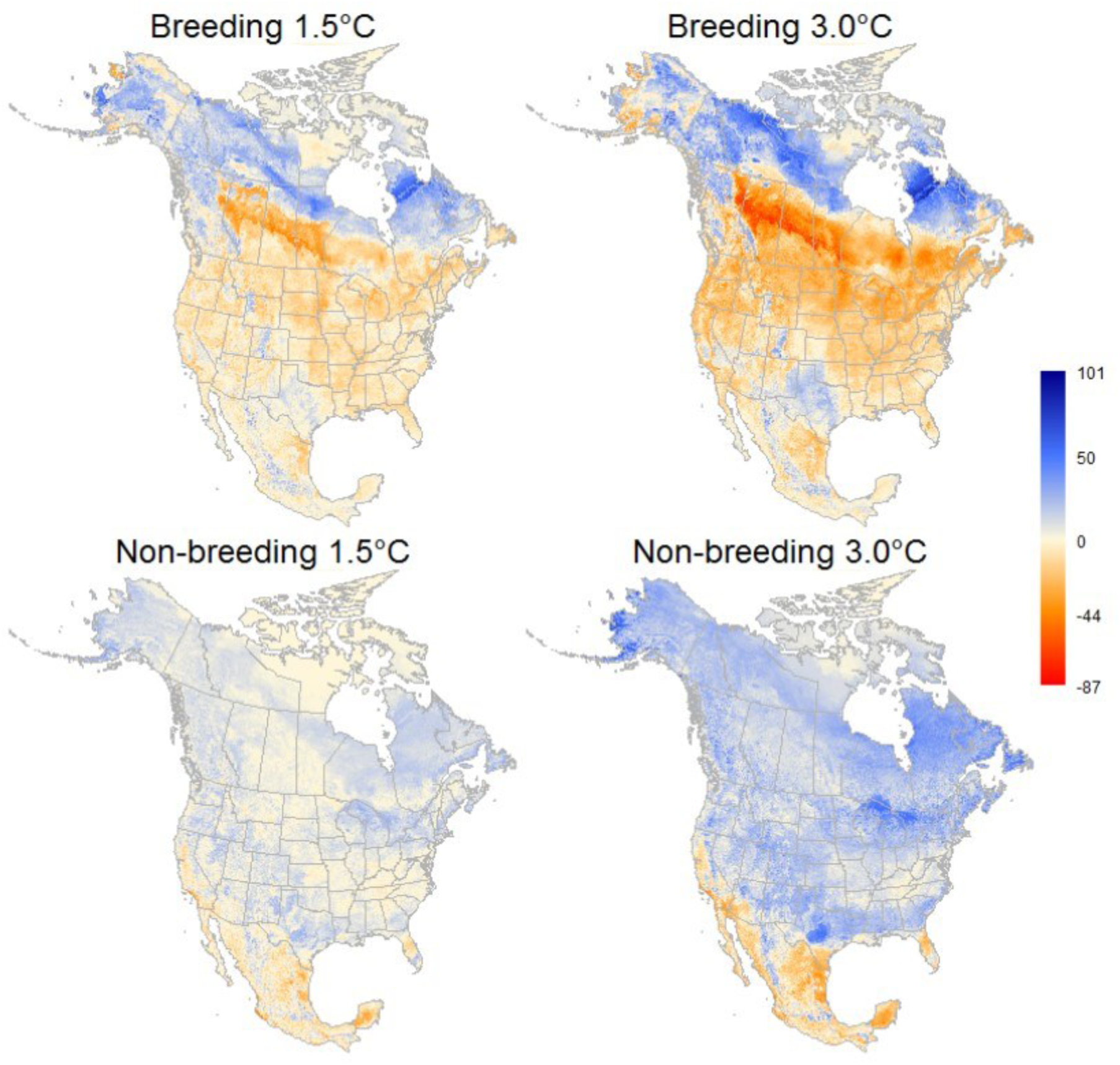
Net gain or loss in number of species at the local community level (i.e. per grid cell) across North America for the breeding and non-breeding seasons. The scale ranges from net loss (red) to a net gain (blue) of species under 1.5°C warming (left) and 3.0°C warming (right). Net change in the breeding season ranged from -87 to 101, and from -62 to 97 in the non-breeding under 3.0°C. Under 1.5°C net change in the breeding seasons ranged from -62 to 82, and from -63 to 58 in the non-breeding.

When looking across bird habitat groups, patterns of net change varied geographically and seasonally (S2 Supplementary Figures and Results, S2.1-S2.12 Figs). Four habitat groups dominated the aggregate patters (Fig 6), including boreal forest (net change min-max: -43 – 40), eastern forest (net change min-max: -26 – 35), waterbirds (net change min-max: -33– 40), and western forests (net change min-max: -35 – 33). The subtropical (S2.9 Fig), boreal (S2.3 Fig) and western forests (S2.12 Fig) groups all have dominant net loss in both seasons, although less pronounced in the non-breeding season (S3.2 Table). Several other groups exhibited loss in the breeding season including arctic (S2.1 Fig), eastern forest (S2.5 Fig), grasslands (S2.7 Fig), and waterbirds S2.11 Fig). Some groups had net gains in both seasons, which included the aridlands, coastal, generalists, marshlands, and urban/suburban groups (S2 Figs, S3.2 Table).

The present conservation status of climate vulnerable species varied widely. Among climate vulnerable species across seasons, 50 were also listed on the PIF Watch List (Fig 7). Thus, the adaptive capacity of these species is further diminished by their current state of conservation need. These include species from the grasslands (7); aridlands (7); coastal (8); sub-tropical (3), western (7), eastern (5), and boreal forests; marshlands (5); waterbirds (5); and arctic (2). Audubon priority species among these include: Baird’s Sparrow, Bobolink, Canada Warbler, Cerulean Warbler, Chestnut-collared Longspur, Florida Scrub Jay, Golden-winged Warbler, Greater Sage-Grouse, Marbled Godwit, Mountain Plover, Piping Plover, Saltmarsh Sparrow, Semipalmated Sandpiper, Sprague’s Pipit, Tricolored Blackbird, and Wood Thrush. Furthermore, 323 of our climate vulnerable species across seasons are not currently on the Watch List (Fig. 7), suggesting and emerging future conservation need as climate change advances.

**Fig. 7.**
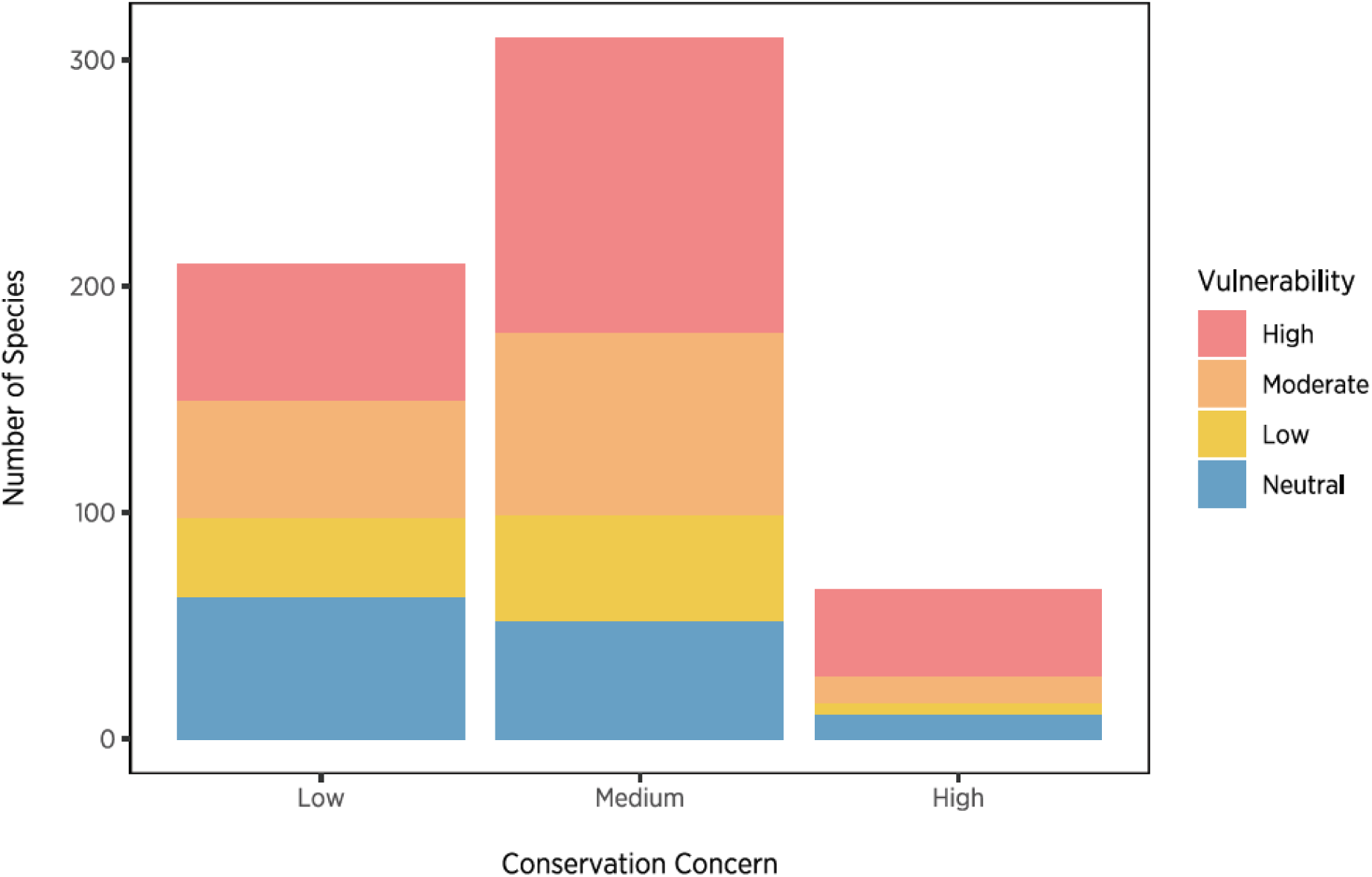
Species vulnerability in the breeding season classified as being present on the PIF Watch list for the 3.0°C scenario across both seasons.

### Model Accuracy

Overall models performed well with cross-validation when predicting to spatially independent datasets (AUC of 0.92 [IQR: 0.87–0.96]) across all species. Between seasons, models performed slightly better in the breeding season, with a median testing AUC of 0.93 (IQR: 0.88–0.97) in breeding and 0.91 (IQR: 0.85–0.95) in non-breeding season. All models across seasons performed better than random with all AUC values above 0.5 and a minimum AUC of 0.62. Maxent (N = 496) out-performed BRT (N = 101) 83% of the time in breeding, and 88% of the time in non-breeding (Maxent, N =483; BRT, N = 63). A spatial resolution of 1 km (breeding N = 386, non-breeding N = 352) outperformed other spatial scales (10 km, breeding N = 146, non-breeding = 148; 25 km, breeding N = 2, non-breeding N = 5; 50 km, breeding N = 63, non-breeding N = 41).

For all species, the median suitability threshold was 0.32 (IQR: 0.24–0.40) with tss (N = 451) being the most common threshold applied, followed by om_10 (N = 338), mk (N = 152), mo (N =123), and eq (N = 79). Across all thresholds, median true positive rate (0.90, IQR: 0.84–0.92) exceeded median true negative rate (0.85, IQR: 0.72–0.93), meaning that, on average, models included 90% of occurrence and excluded 85% of non-occurrence grid cells across species. For true positive and true negative rates by threshold choice, see S2.13 Fig. For species-specific results for AUC, TPR, TNR, threshold choice and value regularization parameters, and model algorithm and spatial filtering resolution selected, See S5 Species Model Evaluation.

In the breeding season, degree-days <0 °C, mean warmest month temperature, and vegetation type were the most important variables across all species (S2.14 Fig). In non-breeding season, mean coldest month temperature, degree-days >5°C, and climatic moisture deficit were the most important variables across all species (S2.14 Fig). In breeding season, temperature variables were most important across all species (median 37.99%), followed closely by precipitation (median 27.88%) and habitat variables (median 12.84%). The same pattern was observed in non-breeding season, although temperature was even more dominant in this season (median 54.25), then precipitation (median 15.91) and habitat (median 9.24). Of the habitat variables, vegetation type was most important across all species (S3.3 Table). Variable importance varied by habitat group and season, with group-specific variables being important for some. Distance to the coast, for example, had a strong influence for coastal species. Other key group variables were surface water occurrence for waterbirds, coastal species, and marshbirds in breeding season and non-breeding season, and the human influence index for urban/suburban species. Distance to shore, wetland type, and distance to wetland had limited influence on models (S3.4 Table).

## DISCUSSION

Our results indicate that nearly two-thirds of North American birds are moderately or highly vulnerable to climate change, confirming that climate change is an emerging threat to birds in North America. Others have found that 53% of North American birds are vulnerable to climate change [36] and that the traits of nearly 24-50% of bird globally make them vulnerable to climate change [4]. Ours is the most comprehensive assessment of climate change vulnerability of bird in North America to date, due to the extensive observation dataset used and to the incorporation of dynamic vegetation and environmental features such as topography, distance to shoreline, and water bodies. However, even this assessment may reasonably be considered conservative because the pace of change is now exceeding even past estimates of climate change [32]. Already in 2017, there was a +1.3°C global land temperature anomaly [73] (NOAA National Centers for Environmental Information [NCEI], 2018), and near 1.0°C average increase already detected globally over pre-industrial levels [32]. Further, the threat of climate change occurs in addition to other threats, such as urbanization and habitat alteration, and land conversion by human activities [74,75].

Climate change mitigation to stabilize warming at 1.5°C globally would reduce vulnerability in North American bird species. These results align with the recommendations of the IPCC, which has identified a target of 2.0°C global temperature increase—preferably 1.5°C—to avoid severe and irreversible effects of climate change [32]. The current pledged reductions in greenhouse gas emissions set in the Paris agreement at present lead to at least 3.2°C global increase in mean temperature [35]. In fact, on our current trajectory, potential warming likely will exceed this (4-5+ °C) by the end of century [76,77] and will result in inevitable warming of 1.5°C globally by 2030-2052. Dramatic and aggressive reductions in emissions that limit warming to 1.5°C would result in 76% of our species classified as vulnerable dropping at least one category in vulnerability, and 38% of those species would be no longer vulnerable. Our results reiterate previous findings that mitigation can have beneficial results, and reductions in emissions by even 0.5°C can greatly reduce the exposure and ramifications of climate change for both human [78] and animal systems [18,79]. This finding is not surprising, given that humans and nature are intertwined and when areas that are important for wildlife (e.g., ecosystem services breakdown) suffer, so do human systems [80]. Our results provide compelling evidence that climate change mitigation that stabilizes warming to less than 3.0°C, and preferably 1.5°C would be beneficial to birds. To do this, we need aggressive climate change mitigation and adequate policies at a national, continental, and global scale.

Current projected change in climate is set to disrupt bird communities across North America. Changes in more than 100 species within a local community are possible under the 3.0°C scenario in both breeding season (dominated by range loss and local extirpation) and non-breeding season (dominated by range gain and local colonization). The reshuffling of species at a continental scale will likely lead to numerous and unpredictable disruptions of ecological function, including loss of key species that could lead to diminished and altered ecosystem health and processes [81]. Changes at this scale could manifest in novel bird assemblages, with communities disassembling and reassembling in new combinations, revealing new species interactions [23]. These novel interactions could lead to population declines and local extinctions if species are not able to adapt quickly enough to new patterns in co-occurrence [82]. With the added complexity of novel communities and ecosystems [83], increased pressure from some invasive species [84], and agricultural intensification [85,86], climate change will be a multi-dimensional, novel experience that will increase species vulnerability, and thus biodiversity loss, unless actions are taken to lessen the effects. Given this, we can use classifications of climate change vulnerability to inform community and species-based climate change adaptation strategies. The most vulnerable species merit the most attention in a triage framework for climate change adaptation [61].

Climate change is an emerging threat to birds in addition to existing threats such as habitat loss, changes in disturbance regimes, and invasive species [87]. Of the 389 climate vulnerable species, 73 are also on the PIF Watchlist. Thus, these species are the priority for conservation triage in a changing climate. However, there were also 422 climate-vulnerable species not include in the PIF watchlist, a pattern replicated with the IUCN Red List, where only 6-9% of the 24-50% of climate change vulnerable bird species globally are currently represented [4]. These species should be the focus of monitoring and observation to determine if the effects of climate change lead to a change in conservation status (Lawler 2009). Furthermore, we identify arctic birds, waterbirds, and boreal and western forest birds to have 78-100% of species vulnerable at 3.0°C warming. Interestingly, those climate-vulnerable habitat groups are not presently classified as being of high conservation concern. These groups merit attention, particularly in higher latitudes where the magnitude of climate change is projected to be greater than global averages [81].

Maps of projected changes in community composition inform place-based climate change adaptation strategies [36,65]. We project large areas of range contraction across species in the northern and eastern temperate forests, the Boreal Shield of Canada, and the northern parts of the Midwest and Northeast in the United States. Canada has already experienced warming at a rate twice as fast relative to the rest of the world, and the arctic has experienced warming three times as fast [88]. This region has a projected shift in vegetation type, with a large contraction of the alpine tundra as arctic tundra and Canadian taiga shift northwards. The alpine and tundra ecoregions are threatened by replacement in the north by Alaskan subarctic conifer forests, and in the south by temperate deciduous forests and Great Plains grasslands [50,89]. Loss of habitat lags in vegetation response, and the pace of climate change in this area makes it vulnerable for birds in the future [89,90]. However, these areas are also considered some of the most important places for breeding birds, deemed the ‘nurseries of North American birds’ where nearly half of all North American species breed [91]. This includes 30% of North America’s landbirds, 30% of shore-birds, and up to 38% of waterbirds.

Our results suggest that range loss will be more pronounced in the breeding season, and outpaces gains across the majority of North America with the exception of the Arctic region. Large losses in breeding season ranges are problematic, given that ranges defined from SDMs have been shown to reflect abundance and carrying capacity, with lower suitability often reflected in lower species abundances [92]. Range loss of this magnitude, therefore, does not bode well for sustainable breeding populations. Disruptions in the breeding season could lead to declines across the species full annual cycle population [93–95]. This idea is proposed by the “tap-hypothesis”, which contends that changes in climate during the breeding season can influence population size for a species by regulating the number of new recruits into the population the following year [96]. Persistence of bird populations is linked to breeding success and productivity, which is often linked to temperature [97–99]. Warming temperatures and changes in climate could potentially reduce breeding productivity, leading to population declines and, in some cases, quasi-extinction when warming is not curbed [93]. In contrast, our results highlight an overall gain in species ranges in the non-breeding season across all scenarios, which have mixed implications. For some species, the increased suitability in the non-breeding season could help balance reduced recruitment in the breeding season through reduced mortality in the non-breeding season-i.e., the “tub-hypothesis” [96], if breeding season loss is minimal. However, if projected species gains in the non-breeding season coincide with large amounts of range loss in the breeding season, then this will be less likely. In some areas, such as the southeastern US, gains are especially pronounced. There appears to be relative stability in climate boundaries for evergreen and deciduous forests [50] in this region, plus a likely influx of range expansions from more southerly species. However, range gains can be problematic too, leading to novel communities and changes in species co-occurrence and interactions [23]. Such large-scale disruptions can even lead to modifications of ecosystem function and net negative consequences for species and ecosystem services [81]. Range gains for species are associated with areas that are newly suitable to a species, indicating that a species would need to shift its range, colonize, and establish new populations. This comes with added complications of species establishment such as competition and multitrophic interactions, as well as specific habitat requirements [100]. Range gains may be problematic for some species, especially when paired with the total loss of historic breeding range. This is seen in Cerulean Warbler (Setophaga cerulean), Black-headed Gull (Chroicocephalus ridibundus), and Golden-cheeked Warbler (Setophaga chrysoparia) all of which are projected to lose more than 90% of their current range, but gain more than 70% elsewhere, or Golden-winged Warbler (Vermivora chrysoptera), which is projected to lose 99% of its current range while gaining 60% in a newly suitable location. For species such as these, it is unknown if range gains in new locations will be sufficient for sustaining populations, given the rapid nature of climate change and the time it takes to establish populations. Lastly, novel climates aren’t projected to peak in the non-breeding season until the first half of the next century [101], suggesting that more changes are to come beyond the temporal extent of our projections.

### Caveats

Our vulnerability assessment of birds in North America is likely a conservative estimate for several reasons. Although our study takes into account future changes in climate and vegetation, we do not assess changes in anthropogenic land use, sea-level rise, extreme events or other global and climate change-related threats to birds. In addition, projected species responses will not always be realized for a variety of reasons, including intrinsic factors such as phenotypic plasticity (e.g., the ability to adapt to changing climate conditions and stay in place), dispersal and colonization abilities, evolutionary potential (i.e., genetic diversity), and extrinsic factors, such as unpredicted habitat loss, species competition, and pollution [102]. We do not assess altered species interactions or how community-level changes could determine species realized future ranges. These were beyond the scope of our current vulnerability assessment, and an identified future research direction. Additionally, we have used a threshold-based approach that only accounts for potential range loss or gains for each species. This accounts for how a species will lose part of its current range, or gain range in new locations (i.e., cross under or over the threshold of suitability), but does not break down changes within a species’ range that are maintained (are currently, and are projected to be above a species threshold of suitability, e.g., ‘stable’). These ‘stable’ areas are in fact changing, with some areas improving or worsening with climate change. Areas that are worsening and approach our selected threshold could indeed be of low suitability, enough to reflect low abundance and population instability [92].

In this work, we strive to balance meeting current standards for SDM-based biodiversity assessments [39] with the effort of modeling 604 species in two seasons. We included an assessment of each species range maps individually by bird exports in both the breeding and non-breeding season, to establish ecological reality of the model outputs. However, we acknowledge that all SDM outputs include inherent uncertainties and errors and therefore do not represent the actual species range. In addition, the use of citizen and community science data sets in the assessment, and the discrepancies of such data collection methods, may violate some assumptions of SDMs [103]. We acknowledge that our bird occurrence data came from various sources, and such were not sampled in a standardized capacity, with uneven sampling due to survey location bias, and with variation in individual detection probabilities and identification skills. We balanced the need to retain as much data as possible to capture species full ranges across North America, with extensive use of spatially structured data partitioning, bias correction, data filtering, and expert review. It has been shown, however, that careful partitioning and analysis of avian data can yield important full life-cycle bird distribution models [104]. Choice of environmental data may also introduce errors and uncertainty, such as the modeled future projections for vegetation data. The vegetation projections used here are not mechanistic and are themselves based on climate models. Climate-based vegetation projections likely overestimate range gains and underestimate vulnerability [89]. Additionally, ensemble vegetation models possibly underestimate vegetation change at 3.0°C and over-estimate change under the 1.5°C scenario [25]. However, continental scale vegetation projections are not readily available, but vegetation provides important information relevant for bird species. Despite these limitations, we chose to include these as the best available variables biologically relevant to this type of modeling. Lastly, uncertainties and errors also arise from the choice of SDM and model selection. Sources of uncertainty are discussed in detail in Wilsey et al. [25].

## CONCLUSION

This analysis is unprecedented in scope in the suite of species modeled and spatial coverage. In addition, it provides a full continental scale assessment of climate change vulnerability of the North American avifauna to inform climate change mitigation and adaptation strategies. Our results provide clear evidence that we need aggressive climate mitigation policies for birds in North America. The dire consequences of a 3.0°C warming scenario, and the reduction in bird species vulnerability if we stabilize global warming at 1.5°C, indicate that we must reduce emissions as much as possible to maintain bird communities. Stabilizing climate at 1.5°C is feasible [33,105], but few deliberate actions have been taken to get us there. The current pledge in greenhouse gas reductions set by the Paris Agreement framework is estimated to equate to a 3.2 °C global increase in mean temperature [35], which is not sufficient to reduce vulnerability to birds.

Birds are early responders to climate change and can be important indicators of large-scale ongoing and future ecological change. Here we also provide clear priorities as to where we could focus adaptation conservation efforts, and for which species we need to alleviate climate change stressors. We identified arctic birds, waterbirds, and boreal and western forest birds as being highly vulnerable to climate change. Interestingly, those habitat groups most threatened by climate change were not classified of high conservation concern presently. In an uncertain future, our study reveals that we must pursue both mitigation and adaptation strategies in order to conserve North American birds in a changing climate. Unless we take aggressive policy mitigation and adaptation actions to alleviate the effects of climate change, birds and the places they need are under threat.

## Supporting information

S2 Supplementary Figure and Results

## ACKNOWLEDGMENTS

This work was funded by the John D. and Catherine T. MacArthur Foundation, grant G-1511-150388 and by U.S. Fish and Wildlife Service funding 140F0318P0263. We also received support from Amazon Web Services’ (AWS) Promotional Credit Program (https://aws.amazon.com/awscredits/) and Domino Data Labs Programs for Non-profits and Education (https://www.dominodatalab.com/domino-for-good/) for data science platform use and cloud processing. The findings and conclusions in this article are those of the author(s) and do not necessarily represent the views of the U.S. Fish and Wildlife Service. We would like to thank all of our partners and the institutions that provided data for this work (listed in the S1 Bird Data Sources) and all of the volunteers and community scientists that contributed to these datasets. We thank our panel of expert reviewers, which included co-author Geoff LeBaron (National Audubon Society), Kenn Kaufman (Kaufman Field Guides), Erik Johnson (Audubon LA), Walker Golder (Audubon NC), Andrea Jones (Audubon CA), Jillian Liner (Audubon NY), Eric Lind (Audubon NY), and Michael Anderson (Ducks Unlimited Canada). We also thank Martha Harbison (National Audubon Society) for improving this manuscript. To the extent possible, this research was completed in compliance with the Guidelines to the Use of Wild Birds in Research; however, the authors were not directly involved in the collection of the datasets used.

## SUPPORTING INFORMATION

**S1.** Bird Data Sources

**S2.** Supplementary Figure and Results

**S3.** Supplementary Tables

## REFERENCES

1. Ceballos G, Ehrlich PR, Barnosky AD, Garcia A, Pringle RM, Palmer TM. Accelerated modern human-induced species losses: Entering the sixth mass extinction. Sci Adv. 2015;1: e1400253–e1400253. doi:10.1126/sciadv.1400253

2. North American Bird Conservation Initiative (NABCI). The State of the Birds 2016. Washington D.C.: U.S. Department of Interior; 2016.

3. Thomas CD, Cameron A, Green RE, Bakkenes M, Beaumont LJ, Collingham YC, et al. Extinction risk from climate change. Nature. 2004;427: 145–8. doi:10.1038/nature02121

4. Foden WB, Butchart SHM, Stuart SN, Vié J-C, Akçakaya HR, Angulo A, et al. Identifying the World’s Most Climate Change Vulnerable Species: A Systematic Trait-Based Assessment of all Birds, Amphibians and Corals. PLoS ONE. 2013;8: e65427. doi:10.1371/journal.pone.0065427

5. Gaston K. The Structure and Dynamics of Geographic Ranges. 2003.

6. Jiménez-Valverde A, Barve N, Lira-Noriega A, Maher SP, Nakazawa Y, Papeş M, et al. Dominant climate influences on North American bird distributions. Glob Ecol Biogeogr. 2011;20: 114–118. doi:10.1111/j.1466-8238.2010.00574.x

7. Root T. Energy Constraints on Avian Distributions and Abundances. Ecology. 1988;69: 330–339. doi:10.2307/1940431

8. White TCR. The role of food, weather and climate in limiting the abundance of animals. Biol Rev. 2008;83: 227–248. doi:10.1111/j.1469-185X.2008.00041.x

9. Bateman BL, Pidgeon AM, Radeloff VC, Allstadt AJ, Akçakaya HR, Thogmartin WE, et al. The importance of range edges for an irruptive species during extreme weather events. Landsc Ecol. 2015;30: 1095–1110. doi:10.1007/s10980-015-0212-6

10. Rahbek C, Graves GR. Multiscale assessment of patterns of avian species richness. Proc Natl Acad Sci. 2001;98: 4534–4539. doi:10.1073/pnas.071034898

11. Rodenhouse NL, Matthews SN, McFarland KP, Lambert JD, Iverson LR, Prasad A, et al. Potential effects of climate change on birds of the Northeast. Mitig Adapt Strateg Glob Change. 2008;13: 517–540. doi:10.1007/s11027-007-9126-1

12. Bellard C, Bertelsmeier C, Leadley P, Thuiller W, Courchamp F. Impacts of climate change on the future of biodiversity. Ecol Lett. 2012;15: 365–377. doi:10.1111/j.1461-0248.2011.01736.x

13. Parmesan C. Ecological and evolutionary responses to recent climate change. Annu Rev Ecol Evol Syst. 2006; 637–669.

14. Hitch AT, Leberg PL. Breeding Distributions of North American Bird Species Moving North as a Result of Climate Change. Conserv Biol. 2007;21: 534–539. doi:10.1111/j.1523-1739.2006.00609.x

15. La Sorte FA, Thompson FR. Poleward shifts in winter ranges of North American birds. Ecology. 2007;88: 1803–1812.

16. Prince K, Zuckerberg B. Climate change in our backyards: the reshuffling of North America’s winter bird communities. Glob Change Biol. 2015;21: 572–585. doi:10.1111/gcb.12740

17. Root TL, Price JT, Hall KR, Schneider SH, Rosenzweig C, Pounds JA. Fingerprints of global warming on wild animals and plants. Nature. 2003;421: 57. doi:10.1038/nature01333

18. Warren R, VanDerWal J, Price J, Welbergen JA, Atkinson I, Ramirez-Villegas J, et al. Quantifying the benefit of early climate change mitigation in avoiding biodiversity loss. Nat Clim Change. 2013;3: 678–682.

19. Wiens JA. Spatial Scaling in Ecology. Funct Ecol. 1989;3: 385–397. doi:10.2307/2389612

20. Mann ME, Zhang Z, Hughes MK, Bradley RS, Miller SK, Rutherford S, et al. Proxy-based reconstructions of hemispheric and global surface temperature variations over the past two millennia. Proc Natl Acad Sci. 2008;105: 13252–13257. doi:10.1073/pnas.0805721105

21. Williams JE, Blois JL. Range shifts in response to past and future climate change: Can climate velocities and species’ dispersal capabilities explain variation in mammalian range shifts? J Biogeogr. 2018;45: 2175–2189. doi:10.1111/jbi.13395

22. Bateman BL, Pidgeon AM, Radeloff VC, VanDerWal J, Thogmartin WE, Vavrus SJ, et al. The pace of past climate change vs. potential bird distributions and land use in the United States. Glob Change Biol. 2016;22: 1130–1144. doi:10.1111/gcb.13154

23. Stralberg D, Jongsomjit D, Howell CA, Snyder MA, Alexander JD, Wiens JA, et al. Re-Shuffling of Species with Climate Disruption: A No-Analog Future for California Birds? PLoS ONE. 2009;4: e6825. doi:10.1371/journal.pone.0006825

24. Williams SE, Shoo LP, Isaac JL, Hoffmann AA, Langham G. Towards an integrated framework for assessing the vulnerability of species to climate change. PLoS Biol. 2008;6: e325.

25. Wilsey C, Taylor L, Brooke Bateman, Jensen C, Michel N, Panjabi A, et al. Climate policy action needed to reduce vulnerability of conservation-reliant grassland birds in North America. Conserv Sci Pract. 2019;CSP2-18–0014.

26. Moritz C, Agudo R. The Future of Species Under Climate Change: Resilience or Decline? Science. 2013;341: 504–508. doi:10.1126/science.1237190

27. Foden WB, Young BE, editors. IUCN SSC Guide-lines for Assessing Species’ Vulnerability to Climate Change [Internet]. IUCN, Cambridge, UK and Gland, Switzerland; 2016. Available: http://www.iucn.org/theme/species/publications/guidelines and http://www.iucn-ccsg.org

28. Morecroft MD, Crick HQP, Duffield SJ, Macgregor NA. Resilience to climate change: translating principles into practice. J Appl Ecol. 2012;49: 547–551. doi:10.1111/j.1365-2664.2012.02136.x

29. IPCC. Climate Change 2007: Impacts, Adaptation and Vulnerability. Contribution of Working Group II to the Fourth Assessment Report of the Intergovernmental Panel on Climate Change. Cambridge: Cambridge University Press; 2007.

30. Wilbanks TJ, Leiby P, Perlack R, Ensminger JT, Wright SB. Toward an integrated analysis of mitigation and adaptation: some preliminary findings. Mitig Adapt Strateg Glob Change. 2007;12: 713–725. doi:10.1007/s11027-007-9095-4

31. IPCC. Summary for Policymakers. In: Stocker TF, Qin D, Plattner G-K, Tignor M, Allen SK, Boschung A, et al., editors. Climate Change 2013: The Physical Science Basis Contribution of Working Group I to the Fifth Assessment Report of the Intergovernmental Panel on Climate Change. Cambridge, United Kingdom and New York, NY, USA: Cambridge University Press; 2013.

32. IPCC. Global warming of 1.5°C [Internet]. Available from http://www.ipcc.ch/report/sr15/; 2018. Available: http://www.ipcc.ch/report/sr15/

33. Meinshausen M, Meinshausen N, Hare W, Raper SCB, Frieler K, Knutti R, et al. Greenhouse-gas emission targets for limiting global warming to 2 °C. Nature. 2009;458: 1158–1162. doi:10.1038/nature08017

34. United Nations. Paris Agreement [Internet]. 2015 [cited 16 Feb 2017]. Available: http://unfccc.int/files/essential_background/convention/application/pdf/english_paris_agreement.pdf

35. Climate Transparency. G20 Brown to Green Report 2018 [Internet]. Climate Transparency; 2018. Available: https://www.climate-transparency.org/g20-climate-performance/g20report2018

36. Langham GM, Schuetz JG, Distler T, Soykan CU, Wilsey C. Conservation Status of North American Birds in the Face of Future Climate Change. PLoS ONE. 2015;10: e0135350. doi:10.1371/journal.pone.0135350

37. Lawler JJ, Shafer SL, White D, Kareiva P, Maurer EP, Blaustein AR, et al. Projected climate-induced faunal change in the Western Hemisphere. Ecology. 2009;90: 588–597. doi:10.1890/08-0823.1

38. Bateman BL, Murphy HT, Reside AE, Mokany K, VanDerWal J. Appropriateness of full-, partial-and no-dispersal scenarios in climate change impact modelling. Divers Distrib. 2013;19: 1224–1234.

39. Araújo MB, Anderson RP, Márcia Barbosa A, Beale CM, Dormann CF, Early R, et al. Standards for distribution models in biodiversity assessments. Sci Adv. 2019;5: eaat4858. doi:10.1126/sciadv.aat4858

40. Wu JX, Wilsey CB, Taylor L, Schuurman GW. Projected avifaunal responses to climate change across the U.S. National Park System. Bosso L, editor. PLOS ONE. 2018;13: e0190557. doi:10.1371/journal.pone.0190557

41. Willis SG, Foden W, Baker DJ, Belle E, Burgess ND, Carr JA, et al. Integrating climate change vulnerability assessments from species distribution models and trait-based approaches. Biol Conserv. 2015;190: 167–178. doi:10.1016/j.biocon.2015.05.001

42. Rodewald P. The Birds of North America [Internet]. 2015. Available: https://birdsna.org

43. Reside AE, Critchell K, Crayn DM, Goosem M, Goosem S, Hoskin CJ, et al. Beyond the model: expert knowledge improves predictions of species’ fates under climate change. Ecol Appl. 2019;29: e01824. doi:10.1002/eap.1824

44. Peterson AT, Soberón J, Pearson RG, Anderson RP, Martínez-Meyer E, Nakamura M, et al. Ecological Niches and Geographic Distributions. Princeton, N.J: Princeton University Press; 2011.

45. Boria RA, Olson LE, Goodman SM, Anderson RP. Spatial filtering to reduce sampling bias can improve the performance of ecological niche models. Ecol Model. 2014;275: 73–77. doi:10.1016/j.ecolmodel.2013.12.012

46. Veloz SD. Spatially autocorrelated sampling falsely inflates measures of accuracy for presence-only niche models. J Biogeogr. 2009;36: 2290–2299.

47. Phillips SJ, Dudík M, Elith J, Graham CH, Lehmann A, Leathwick J, et al. Sample selection bias and presence-only distribution models: implications for background and pseudo-absence data. Ecol Appl. 2009;19: 181–197.

48. Wang T, Hamann A, Spittlehouse D, Carroll C. Locally Downscaled and Spatially Customizable Climate Data for Historical and Future Periods for North America. PLOS ONE. 2016;11: e0156720. doi:10.1371/journal.pone.0156720

49. Bateman BL, Pidgeon AM, Radeloff VC, Flather CH, VanDerWal J, Akçakaya HR, et al. Potential breeding distributions of U.S. birds predicted with both short-term variability and long-term average climate data. Ecol Appl. 2016;26: 2720–2731. doi:10.1002/eap.1416

50. Rehfeldt GE, Crookston NL, Sáenz-Romero C, Campbell EM. North American vegetation model for land-use planning in a changing climate: a solution to large classification problems. Ecol Appl. 2012;22: 119–141. doi:10.1890/11-0495.1

51. Riley S, Degloria S, Elliot SD. A Terrain Ruggedness Index that Quantifies Topographic Heterogeneity. Int J Sci. 1999;5: 23–27.

52. Canada Centre for Remote Sensing (CCRS), Comisión Nacional para el Conocimiento y Uso de la Biodiversidad (CONABIO), Comisión Nacional Forestal (CONAFOR), Insituto Nacional de Estadística y Geografía (INEGI), U.S. Geological Survey (USGS). 2010 Land Cover of North America at 250 meters [Internet]. Montréal, Québec, Canada: Commission for Environmental Cooperation; 2013. Available: http://www.cec.org/naatlas/

53. Pekel J-F, Cottam A, Gorelick N, Belward AS. High -resolution mapping of global surface water and its long-term changes. Nature. 2016;540: 418–422. doi:10.1038/nature20584

54. Lehner B, Döll P. Development and validation of a global database of lakes, reservoirs and wetlands. J Hydrol. 2004;296: 1–22.

55. Wessel P, Smith WHF. A global, self-consistent, hierarchical, high-resolution shoreline database. J Geophys Res Solid Earth. 1996;101: 8741–8743. doi:10.1029/96JB00104

56. Wildlife Conservation Society-WCS; Center For International Earth Science Information Network-CIESIN-Columbia University. Last of the Wild Project, Version 2, 2005 (LWP-2): Global Human Influence Index (HII) Dataset (Geographic) [Internet]. Palisades, NY: NASA Socioeconomic Data and Applications Center (SEDAC); 2005. doi:10.7927/h4bp00qc

57. Knutti R, Sedláček J. Robustness and uncertainties in the new CMIP5 climate model projections. Nat Clim Change. 2013;3: 369–373. doi:10.1038/nclimate1716

58. GDAL/OGR contributors. GDAL/OGR Geospatial Data Abstraction Library [Internet]. Open Source Geospatial Foundation; 2018. Available: http://gdal.org

59. McGill BJ. Matters of Scale. Science. 2010;328: 575–576. doi:10.1126/science.1188528

60. Seo C, Thorne JH, Hannah L, Thuiller W. Scale effects in species distribution models: implications for conservation planning under climate change. Biol Lett. 2008; Available: https://royalsocietypublishing.org/doi/abs/10.1098/rsbl.2008.0476

61. Lawler JJ. Climate change adaptation strategies for resource management and conservation planning. Ann N Y Acad Sci. 2009;1162: 79–98.

62. Radosavljevic A, Anderson RP. Making better Maxent models of species distributions: complexity, overfitting and evaluation. J Biogeogr. 2014;41: 629–643.

63. Wenger SJ, Olden JD. Assessing transferability of ecological models: an underappreciated aspect of statistical validation. Methods Ecol Evol. 2012;3: 260–267. doi:10.1111/j.2041-210X.2011.00170.x

64. Wilsey CB, Jensen CM, Miller N. Quantifying avian relative abundance and ecosystem service value to identify conservation opportunities in the Midwestern U.S. Avian Conserv Ecol. 2016;11. doi:10.5751/ACE-00902-110207

65. Elith J, Leathwick J. Boosted Regression Trees for ecological modeling [Internet]. 2014. Available: http://www.idg.pl/mirrors/CRAN/web/packages/dismo/vignettes/brt.pdf

66. VanDerWal J, Falconi L, Januchowski S, Shoo L, Storlie C. SDMTools: Species distributions modeling tools: tools for precessing data associated with species distribution modeling exercises. R package version 1.1-221. HttpsCRANR-Proj. 2014;

67. BirdLife International. IUCN Red List for birds [Internet]. 2017 [cited 1 Jul 2016]. Available: http://www.birdlife.org

68. Beauchamp G. Group-foraging is not associated with longevity in North American birds. Biol Lett. 2009; rsbl20090691. doi:10.1098/rsbl.2009.0691

69. Thomas CD, Hill JK, Anderson BJ, Bailey S, Beale CM, Bradbury RB, et al. A framework for assessing threats and benefits to species responding to climate change. Methods Ecol Evol. 2010;2: 125–142. doi:10.1111/j.2041-210X.2010.00065.x

70. Rosenberg KV, Kennedy JA, Dettmers R, Ford RP, Reynolds D, Alexander JD, et al. Partners in flight landbird conservation plan: 2016 revision for Canada and continental United States. Partners in Flight Science Committee; 2016 p. 119.

71. Hole DG, Willis SG, Pain DJ, Fishpool LD, Butchart SHM, Collingham YC, et al. Projected impacts of climate change on a continent-wide protected area network. Ecol Lett. 2009;12: 420–431. doi:10.1111/j.1461-0248.2009.01297.x

72. Hole DG, Huntley B, Arinaitwe J, Butchart SHM, Collingham YC, Fishpool LDC, et al. Toward a Management Framework for Networks of Protected Areas in the Face of Climate Change: Management of Protected-Area Networks. Conserv Biol. 2011;25: 305–315. doi:10.1111/j.1523-1739.2010.01633.x

73. NOAA National Centers for Environmental Information (NCEI). State of the Climate: Global Climate Report - Annual 2017 [Internet]. 2018 [cited 2 Jul 2018]. Available: https://www.ncdc.noaa.gov/sotc/global/201713

74. Radeloff VC, Hammer RB, Stewart SI, Fried JS, Holcomb SS, McKeefry JF. The Wildland–Urban Interface in the United States. Ecol Appl. 2005;15: 799–805. doi:10.1890/04-1413

75. Radeloff VC, Nelson E, Plantinga AJ, Lewis DJ, Helmers D, Lawler JJ, et al. Economic-based projections of future land use in the conterminous United States under alternative policy scenarios. Ecol Appl. 2012;22: 1036–1049. doi:10.1890/11-0306.1

76. Collins M, Knutti R, Arblaster J, Dufresne J-L, Fichefet T, Friedlingstein P, et al. Chapter 12 - Long-term climate change: Projections, commitments and irreversibility. In: IPCC, editor. Climate Change 2013: The Physical Science Basis IPCC Working Group I Contribution to AR5. Cambridge: Cambridge University Press; 2013. Available: http://www.climatechange2013.org/images/report/WG1AR5_Chapter12_FINAL.pdf

77. New Mark, Liverman Diana, Schroder Heike, Anderson Kevin. Four degrees and beyond: the potential for a global temperature increase of four degrees and its implications. Philos Trans R Soc Math Phys Eng Sci. 2011;369: 6–19. doi:10.1098/rsta.2010.0303

78. Shindell D, Faluvegi G, Seltzer K, Shindell C. Quantified, localized health benefits of accelerated carbon dioxide emissions reductions. Nat Clim Change. 2018;8: 291. doi:10.1038/s41558-018-0108-y

79. Warren R, Price J, Graham E, Forstenhaeusler N, VanDerWal J. The projected effect on insects, vertebrates, and plants of limiting global warming to 1.5°C rather than 2°C. Science. 2018;360: 791–795. doi:10.1126/science.aar3646

80. Steffen W, Persson Å, Deutsch L, Zalasiewicz J, Williams M, Richardson K, et al. The Anthropocene: From Global Change to Planetary Stewardship. Ambio. 2011;40: 739–761. doi:10.1007/s13280-011-0185-x

81. Barbet-Massin M, Jetz W. The effect of range changes on the functional turnover, structure and diversity of bird assemblages under future climate scenarios. Glob Change Biol. 2015;21: 2917–2928. doi:10.1111/gcb.12905

82. Hughes L. Biological consequences of global warming: is the signal already apparent? Trends Ecol Evol. 2000;15: 56–61. doi:10.1016/S0169-5347(99)01764-4

83. Radeloff VC, Williams JW, Bateman BL, Burke KD, Carter SK, Childress ES, et al. The rise of novelty in ecosystems. Ecol Appl. 2015;25: 2051–2068. doi:10.1890/14-1781.1

84. Bellard C, Thuiller W, Leroy B, Genovesi P, Bakkenes M, Courchamp F. Will climate change promote future invasions? Glob Change Biol. 2013;19: 3740–3748. doi:10.1111/gcb.12344

85. Tscharntke T, Clough Y, Wanger TC, Jackson L, Motzke I, Perfecto I, et al. Global food security, biodiversity conservation and the future of agricultural intensification. Biol Conserv. 2012;151: 53–59. doi:10.1016/j.biocon.2012.01.068

86. Stanton RL, Morrissey CA, Clark RG. Analysis of trends and agricultural drivers of farmland bird declines in North America: A review. Agric Ecosyst Environ. 2018;254: 244–254. doi:10.1016/j.agee.2017.11.028

87. Wade AA, Hand BK, Kovach RP, Muhlfeld CC, Waples RS, Luikart G. Assessments of species’ vulnerability to climate change: from pseudo to science. Biodivers Conserv. 2016; 1–7. doi:10.1007/s10531-016-1232-5

88. Bush E, Lemmen DS. Canada’s Changing Climate Report [Internet]. Ottawa, ON.: Government of Canada; 2019 p. 444. Available: https://changingclimate.ca/CCCR2019/about/

89. Stralberg D, Bayne EM, Cumming SG, Sólymos P, Song SJ, Schmiegelow FKA. Conservation of future boreal forest bird communities considering lags in vegetation response to climate change: a modified refugia approach. Loyola R, editor. Divers Distrib. 2015;21: 1112–1128. doi:10.1111/ddi.12356

90. Cadieux P, Boulanger Y, Cyr D, Taylor AR, Price DT, Tremblay JA. Spatially explicit climate change projections for the recovery planning of threatened species: The Bicknell’s Thrush (Catharus Bicknelli) as a case study. Glob Ecol Conserv. 2019;17: e00530. doi:10.1016/j.gecco.2019.e00530

91. Blancher P, Wells J. The Boreal Forest Region: North America’s Bird Nursery. Bird Studies Canada and Boreal Songbird Initiative; 2005.

92. VanDerWal J, Shoo LP, Johnson CN, Williams SE. Abundance and the Environmental Niche: Environmental Suitability Estimated from Niche Models Predicts the Upper Limit of Local Abundance. Am Nat. 2009;174: 282–291. doi:10.1086/600087

93. Bonnot TW, Cox WA, Thompson FR, Millspaugh JJ. Threat of climate change on a songbird population through its impacts on breeding. Nat Clim Change. 2018;8: 718. doi:10.1038/s41558-018-0232-8

94. Crick HQP. The impact of climate change on birds. Ibis. 2004;146: 48–56. doi:10.1111/j.1474-919X.2004.00327.x

95. Sæther B-E, Engen S. Population consequences of climate change. Effects of Climate Change on Birds. New York: Oxford University Press; 2010. p. 321.

96. SÆther B-E, Sutherland WJ, Engen S. Climate Influences on Avian Population Dynamics. Advances in Ecological Research. Academic Press; 2004. pp. 185–209. doi:10.1016/S0065-2504(04)35009-9

97. Cox WA, Thompson FR, Reidy JL. The Effects of Temperature on Nest Predation by Mammals, Birds, and Snakes. Efectos de la Temperatura en la Depredación de nidos por parte de Mamíferos, Aves y Serpientes. Auk Ornithol Adv. 2013;130: 784–790. doi:10.1525/auk.2013.13033

98. Cox WA, Thompson FR, Reidy JL, Faaborg J. Temperature can interact with landscape factors to affect songbird productivity. Glob Change Biol. 2013;19: 1064–1074. doi:10.1111/gcb.12117

99. Townsend AK, Cooch EG, Sillett TS, Rodenhouse NL, Holmes RT, Webster MS. The interacting effects of food, spring temperature, and global climate cycles on population dynamics of a migratory songbird. Glob Change Biol. 2016;22: 544–555. doi:10.1111/gcb.13053

100. Van der Putten WH, Macel M, Visser ME. Predicting species distribution and abundance responses to climate change: why it is essential to include biotic interactions across trophic levels. Philos Trans R Soc B Biol Sci. 2010;365: 2025–2034. doi:10.1098/rstb.2010.0037

101. La Sorte FA, Fink D, Johnston A. Time of emergence of novel climates for North American migratory bird populations. Ecography. 2019;0. doi:10.1111/ecog.04408

102. Beever EA, O’Leary J, Mengelt C, West JM, Julius S, Green N, et al. Improving conservation outcomes with a new paradigm for understanding species’ fundamental and realized adaptive capacity. Conserv Lett. 2015; Available: http://onlinelibrary.wiley.com/doi/10.1111/conl.12190/pdf

103. Yackulic CB, Chandler R, Zipkin EF, Royle JA, Nichols JD, Grant EHC, et al. Presence-only modelling using MAXENT: when can we trust the inferences? Methods Ecol Evol. 2013;4: 236–243. doi:10.1111/2041-210x.12004

104. Hochachka WM, Fink D, Hutchinson RA, Sheldon D, Wong W-K, Kelling S. Data-intensive science applied to broad-scale citizen science. Trends Ecol Evol. 2012;27: 130–137. doi:10.1016/j.tree.2011.11.006

105. Goodwin P, Brown S, Haigh ID, Nicholls RJ, Matter JM. Adjusting Mitigation Pathways to Stabilize Climate at 1.5° C and 2.0°C Rise in Global Temperatures to Year 2300. Earths Future. 2018;6: 601–615. doi:10.1002/2017EF000732

