## Supplementary material for "North American Birds Require Mitigation and Adaptation to Reduce Vulnerability to Climate Change": S2 Supplementary Figure and Results

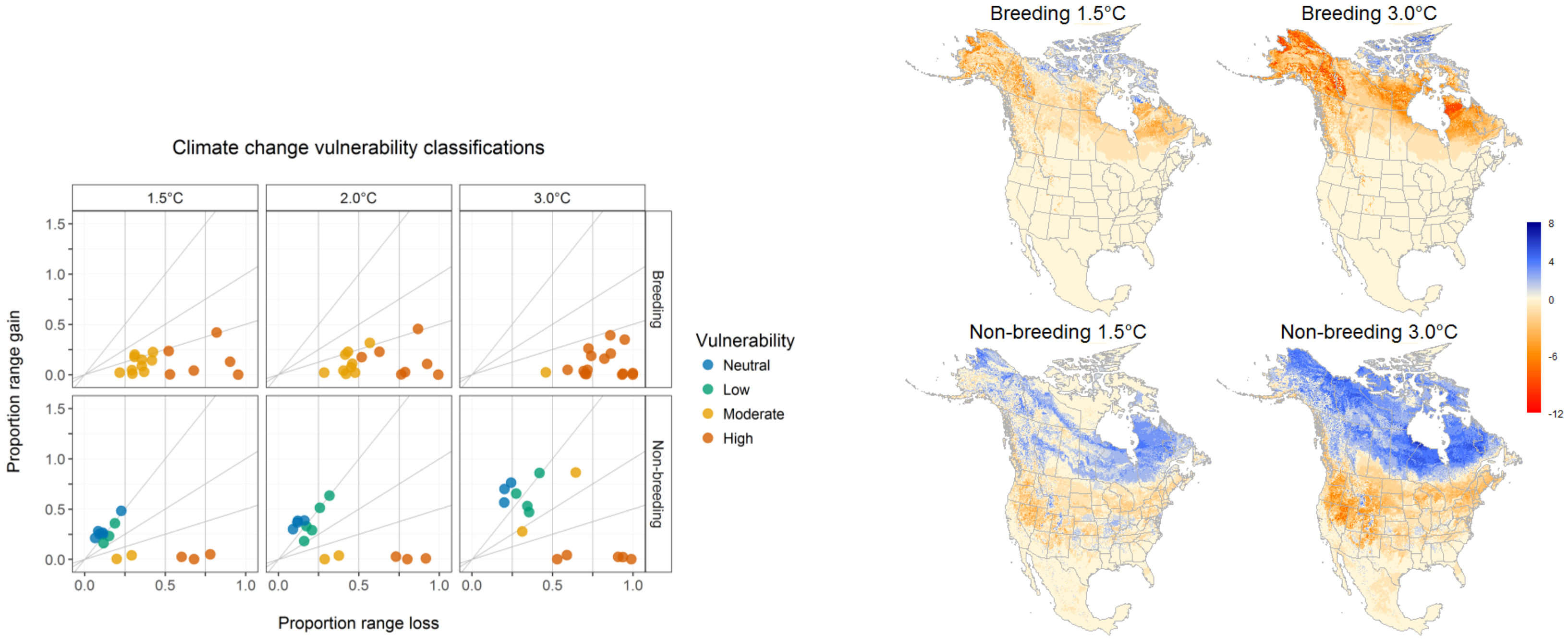


S2.1 Fig. Arctic bird species vulnerability plot and net change maps.

Arctic bird species results for a) vulnerability of all Arctic species in breeding and non-breeding seasons under 1.5 °C, 2.0 °C and 3.0 °C global warming scenarios, and b) Net gain or loss in number of species at the local community level across Arctic species for the breeding and non-breeding seasons. The scale ranges from net loss (red) to a net gain (blue) of species under 1.5°C warming (left) and 3.0°C warming (right). Net change in the breeding season range from -12 to 7, and from -9 to 7 species in the non-breeding under 3.0°C. Under 1.5°C net change in the breeding seasons ranges from -10 to 8, and from -6 to 6 species in the non-breeding. The Arctic group had the highest median range loss in the breeding season, with 74% median range loss spread out across all of Alaska and most of Canada except the northern most part of the tundra and arctic regions (S3.2 Table). In the non-breeding season, we see strong gains with the arctic group across this same region, but was paired with loss across most of the lower 48 states.

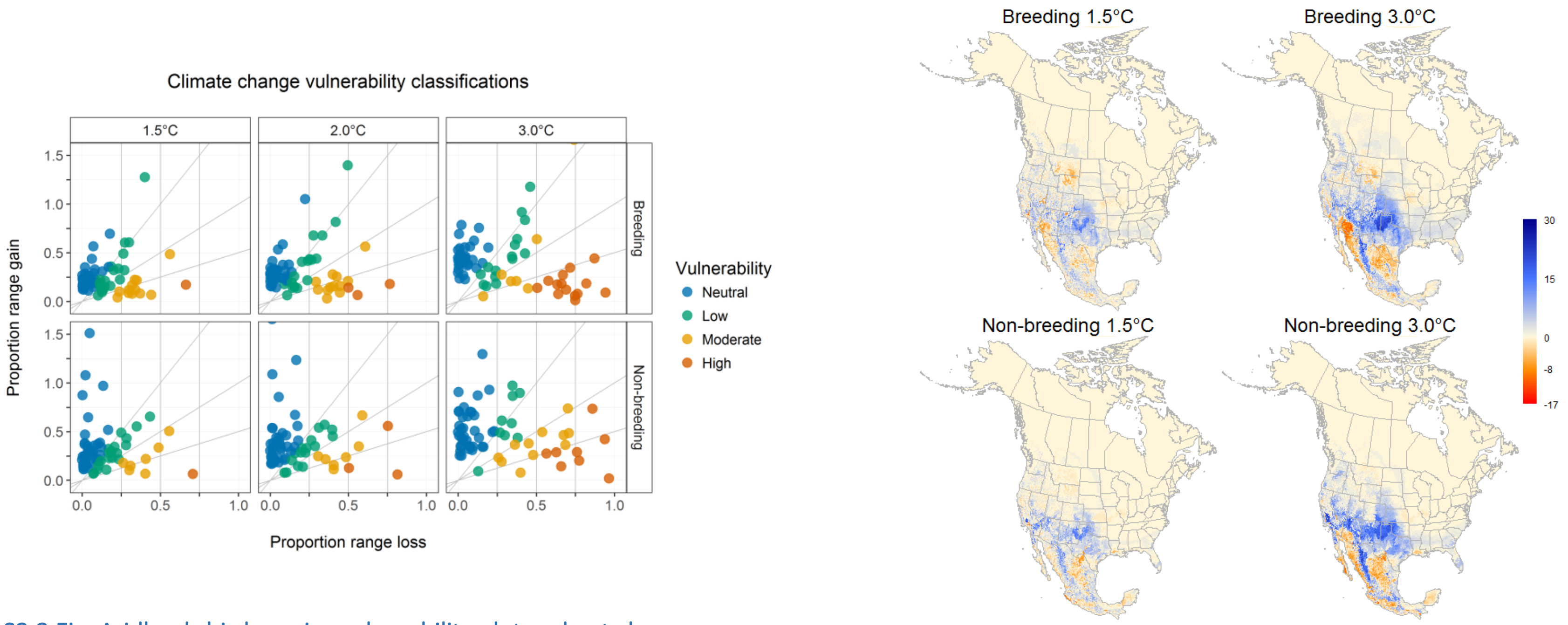


S2.2 Fig. Aridlands bird species vulnerability plot and net change maps.

Aridlands bird species results for a) vulnerability of all Aridlands species in breeding and non-breeding seasons under 1.5 °C, 2.0 °C and 3.0 °C global warming scenarios, and b) Net gain or loss in number of species at the local community level across Aridlands species for the breeding and non-breeding seasons. The scale ranges from net loss (red) to a net gain (blue) of species under 1.5°C warming (left) and 3.0°C warming (right). Net change in the breeding season range from -15 to 27, and from -18 to 30 species in the non-breeding under 3.0°C. Under 1.5°C net change in the breeding seasons ranges from -14 to 21, and from -17 to 26 species in the non-breeding. Across both seasons, the Aridlands group gains outpace loss (median net range gain of 19.5% in breeding and 32.6% in non-breeding, S3.2 Table), especially in the lower Great Plains and North American Deserts with range expansion northward.

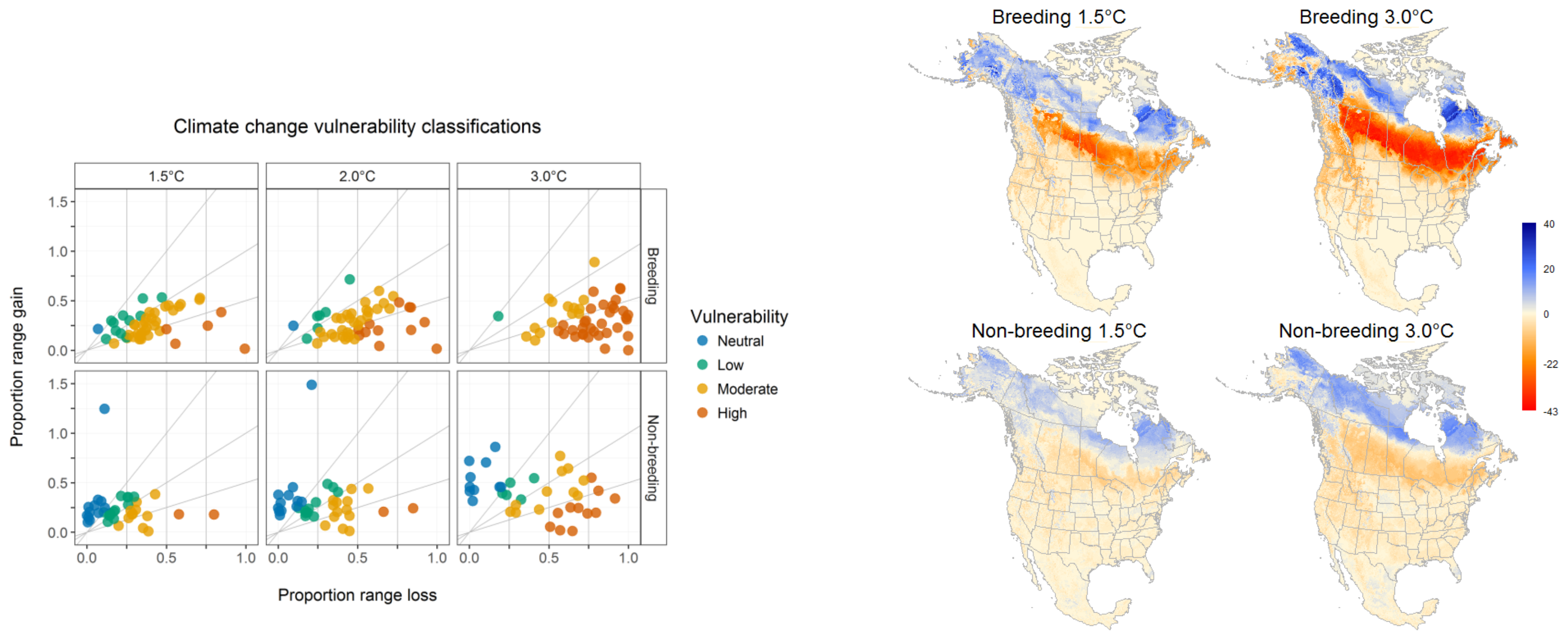


S2.3 Fig. Boreal forests bird species vulnerability plot and net change maps.

Boreal forest bird species results for a) vulnerability of all Boreal forest species in breeding and non-breeding seasons under 1.5 °C, 2.0 °C and 3.0 °C global warming scenarios, and b) Net gain or loss in number of species at the local community level across Boreal forest species for the breeding and non-breeding seasons. The scale ranges from net loss (red) to a net gain (blue) of species under 1.5°C warming (left) and 3.0°C warming (right). Net change in the breeding season range from -43 to 40, and from -14 to 19 species in the non-breeding under 3.0°C. Under 1.5°C net change in the breeding seasons ranges from -35 to 38, and from -11 to 18 species in the non-breeding. The Boreal forests group had 45.1% median net range loss in breeding and near equal rates of loss and gain in the non-breeding; in both seasons, loss was dominant across the boreal Northern Forests, taiga and Hudson Plain, with gains seen in the taiga, tundra and arctic (S3.2 Table).

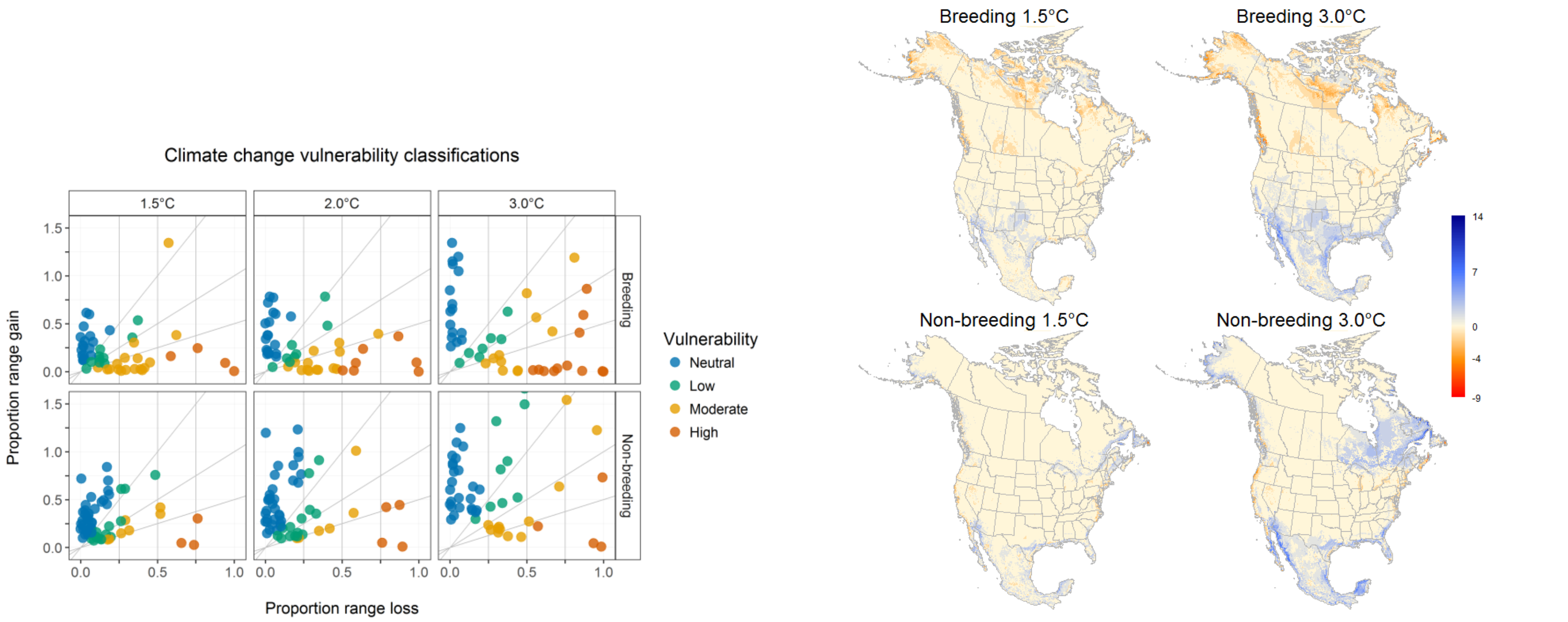


S2.4 Fig. Coastal bird species vulnerability plot and net change maps.

Coastal bird species results for a) vulnerability of all Coastal species in breeding and non-breeding seasons under 1.5 °C, 2.0 °C and 3.0 °C global warming scenarios, and b) Net gain or loss in number of species at the local community level across Coastal species for the breeding and non-breeding seasons. The scale ranges from net loss (red) to a net gain (blue) of species under 1.5°C warming (left) and 3.0°C warming (right). Net change in the breeding season range from -9 to 10, and from -9 to 14 species in the non-breeding under 3.0°C. Under 1.5°C net change in the breeding seasons ranges from -9 to 8, and from -9 to 13 species in the non-breeding. Cells in Figure S2.4 are aggregated to a 10-km resolution to improve visualization along a narrow coastline, but ranges are provided based on original 1-km data values. In the Coastal group, gains are more pronounce din the non-breeding season (median net range gain of 35.2%), than in the breeding season (median net range gain of 6.3%). In both seasons, we see gains across much of Mexico, Florida, and the Southeast (S3.2 Table). In the breeding season, we see range loss across most of coastal Canada and Alaska, and in the non-breeding season along the west and northeast and northwest coasts of the USA.

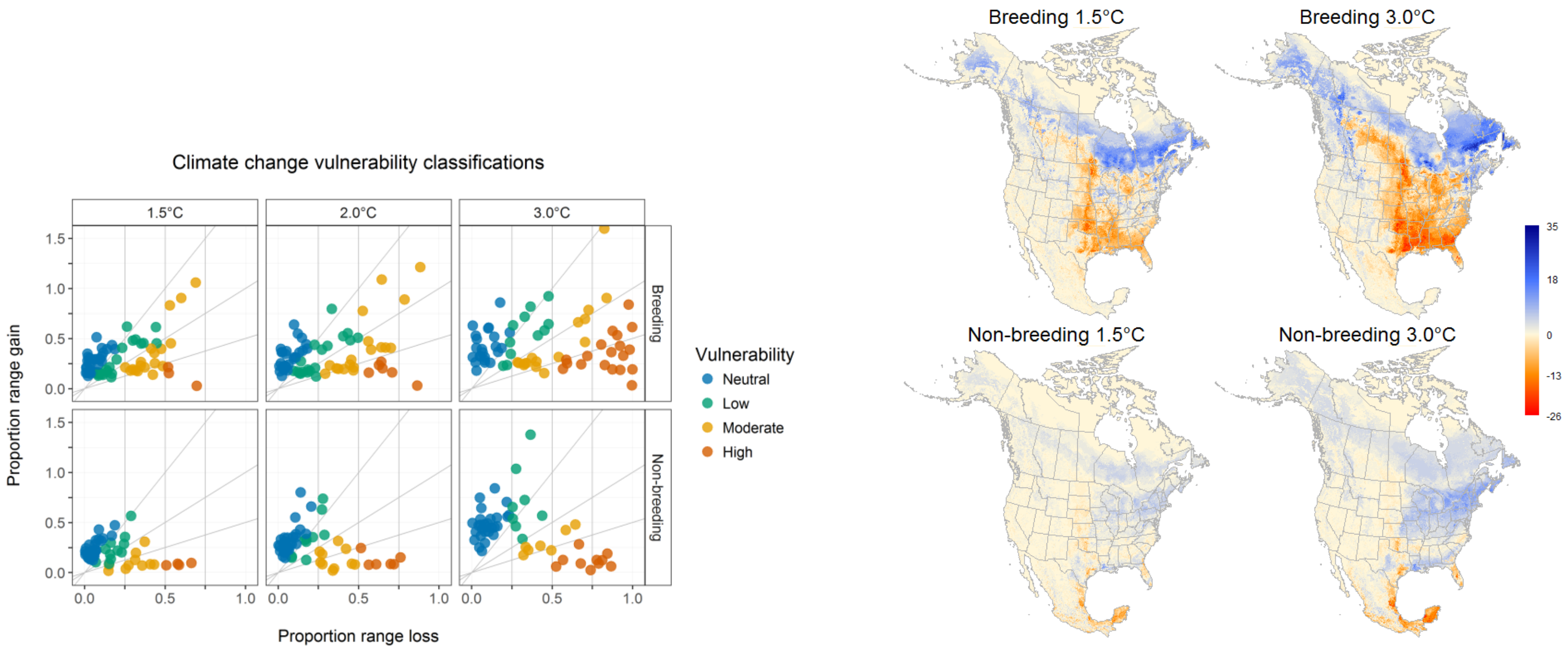


### S2.5 Fig. Eastern Forests bird species vulnerability plot and net change maps.

Eastern forest bird species results for a) vulnerability of all Eastern forest species in breeding and non-breeding seasons under 1.5 °C, 2.0 °C and 3.0 °C global warming scenarios, and b) Net gain or loss in number of species at the local community level across Eastern forest species for the breeding and non-breeding seasons. The scale ranges from net loss (red) to a net gain (blue) of species under 1.5°C warming (left) and 3.0°C warming (right). Net change in the breeding season range from -26 to 35, and from -23 to 18 species in the non-breeding under 3.0°C. Under 1.5°C net change in the breeding seasons ranges from -22 to 28, and from -19 to 16 species in the non-breeding. For the Eastern forests (S3.2 Table), loss slightly outpaced gains in breeding (-0.4% median net loss), but gains dominate in the non-breeding season. The focus in range loss in the breeding season is in the tropical wet forests and eastern temperate forests and Boreal Plain of the Northern forests.  In the non-breeding season, we see range gains in similar areas to breeding season loss with the exception of the tropical wet forests were loss is still dominant.

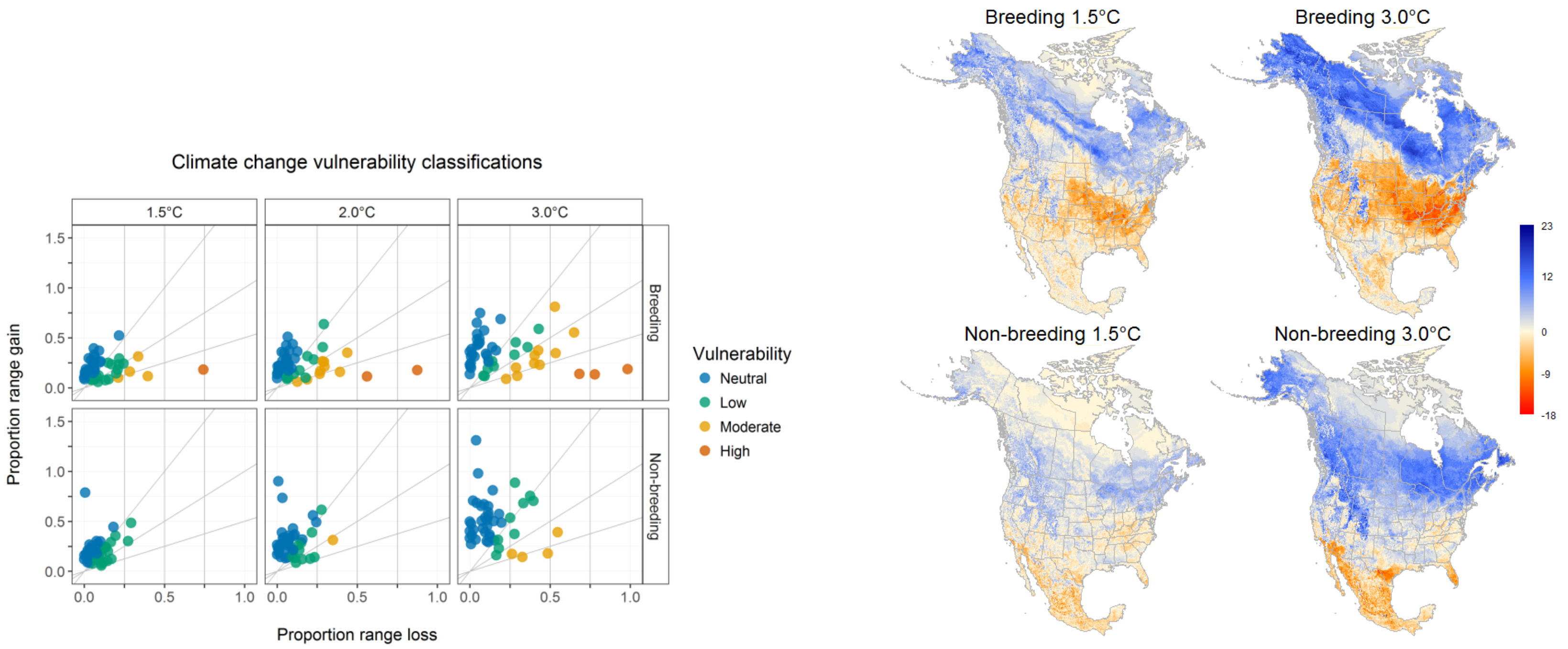


### S2.6 Fig. Generalists bird species vulnerability plot and net change maps.

Generalist bird species results for a) vulnerability of all Generalist species in breeding and non-breeding seasons under 1.5 °C, 2.0 °C and 3.0 °C global warming scenarios, and b) Net gain or loss in number of species at the local community level across Generalist species for the breeding and non-breeding seasons. The scale ranges from net loss (red) to a net gain (blue) of species under 1.5°C warming (left) and 3.0°C warming (right). Net change in the breeding season range from -18 to 23, and from -16 to 20 species in the non-breeding under 3.0°C. Under 1.5°C net change in the breeding seasons ranges from -14 to 17, and from -16 to 12 species in the non-breeding. For Generalist bird species, we see gains in both the breeding (median net range gain 21.2%) and non-breeding (median net range gain 36.3%) broadly across Canada and Alaska (S3.2 Table). Loss is more wide-spread in the breeding season, concentrated in Mexico in Florida in the non-breeding and expanding northward to most of the lower 48 with the exception of the northwestern forests mountains.

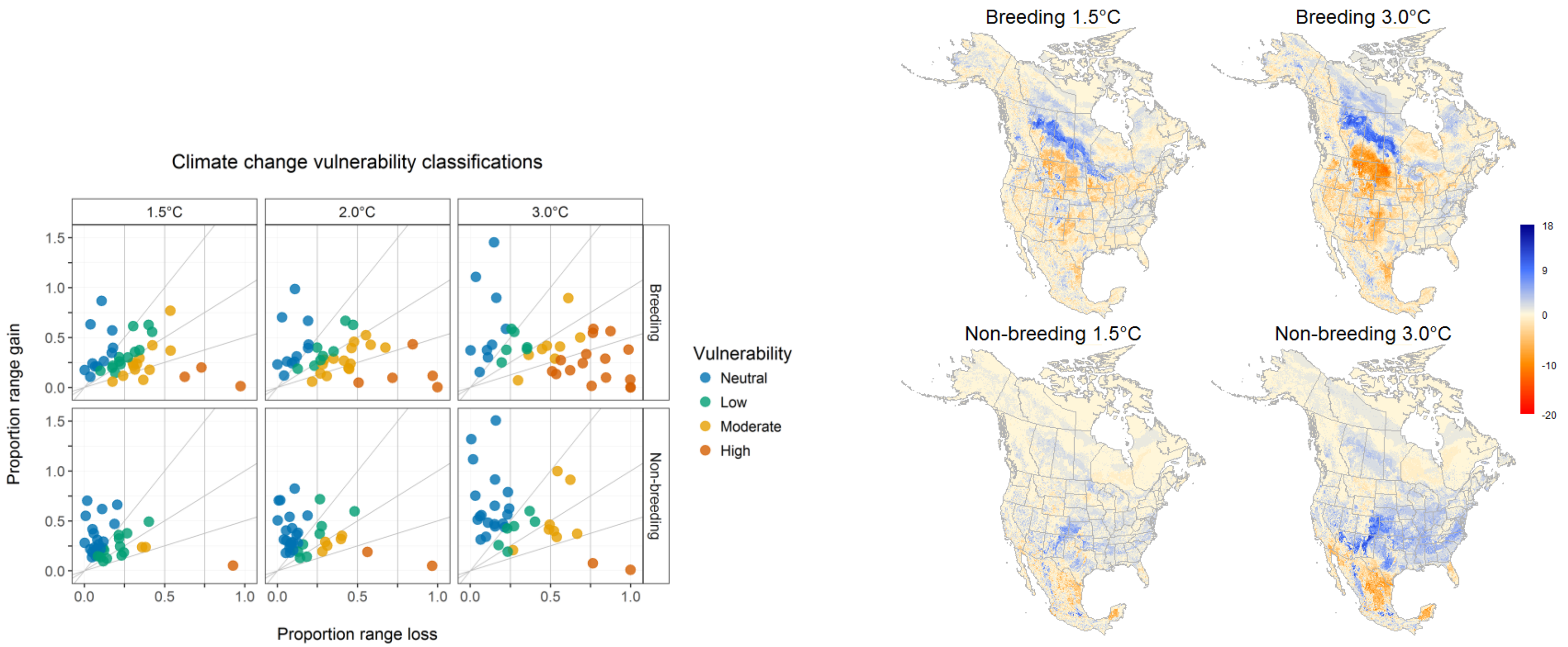


### S2.7 Fig. Grasslands bird species vulnerability plot and net change maps.

Grassland bird species results for a) vulnerability of all Grassland species in breeding and non-breeding seasons under 1.5 °C, 2.0 °C and 3.0 °C global warming scenarios, and b) Net gain or loss in number of species at the local community level across Grassland species for the breeding and non-breeding seasons. The scale ranges from net loss (red) to a net gain (blue) of species under 1.5°C warming (left) and 3.0°C warming (right). Net change in the breeding season range from -15 to 14, and from -20 to 16 species in the non-breeding under 3.0°C. Under 1.5°C net change in the breeding seasons ranges from -15 to 18, and from -18 to 14 species in the non-breeding. In Grassland bird species, -13.9% median net range loss in the breeding season were focused in the Great Plains region, with range gains shifting northward into the Northern forests and taiga (S3.2 Table). For this group, range gains outpaced loss in winter, with most gains concentrated in the eastern temperate forests region, although we see loss in Mexico and Florida as species ranges shift northward.

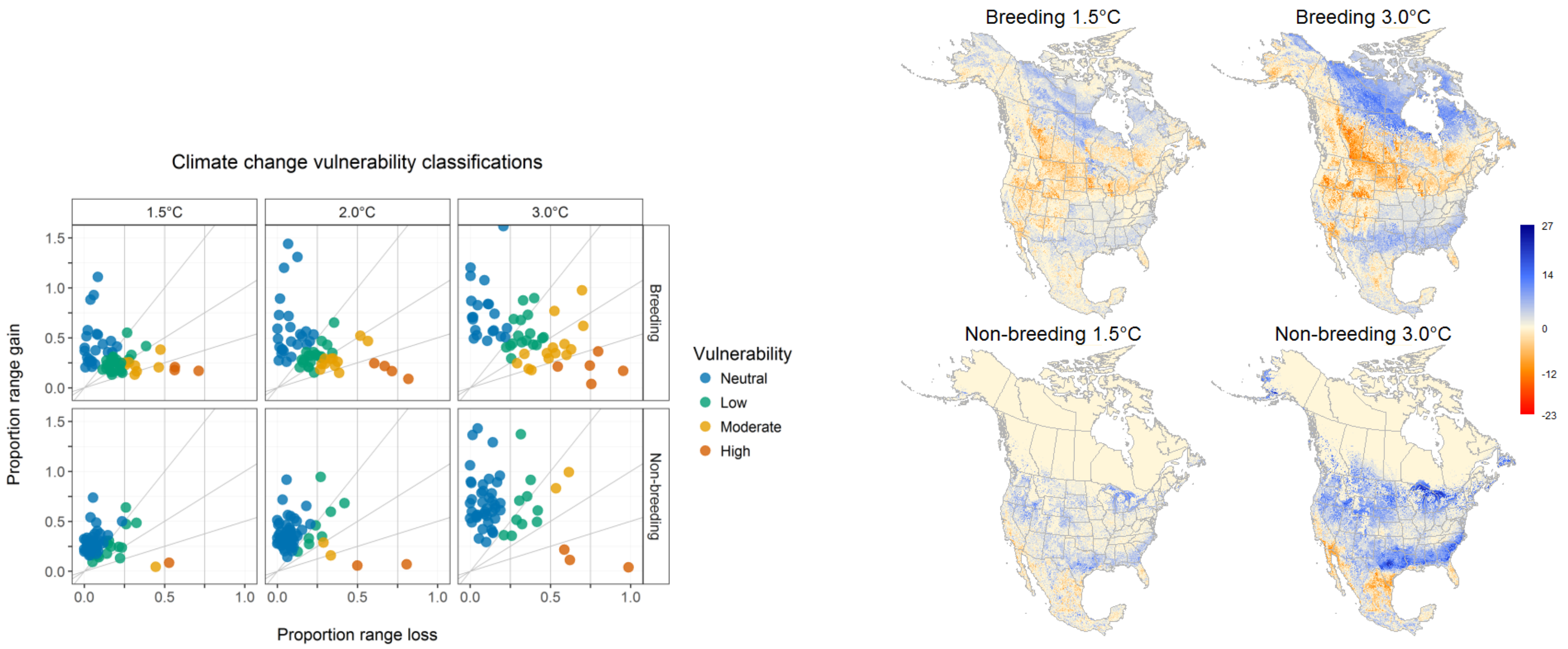


S2.8 Fig. Marshlands bird species vulnerability plot and net change maps.

Marshland bird species results for a) vulnerability of all Marshland species in breeding and non-breeding seasons under 1.5 °C, 2.0 °C and 3.0 °C global warming scenarios, and b) Net gain or loss in number of species at the local community level across marshland species for the breeding and non-breeding seasons. The scale ranges from net loss (red) to a net gain (blue) of species under 1.5°C warming (left) and 3.0°C warming (right). Net change in the breeding season range from -23 to 24, and from -19 to 27 species in the non-breeding under 3.0°C. Under 1.5°C net change in the breeding seasons ranges from -21 to 20, and from -14 to 19 species in the non-breeding. Our Marshlands group saw strong net gain in both the breeding (median range gain of 21.95%) and non-breeding (median net range gain of 47.7%) seasons (S3.2 Table). For this group, breeding and non-breeding season gains differed, with gains seen in the southeastern temperate forests, taiga and tundra regions in the breeding season with loss across most of the rest of the US and Canada. In the non-breeding season, we see gains across most of the lower 48, and loss concentrated in Florida, the Great Plains of southern Texas and Mexico, and the Mexican North American deserts.

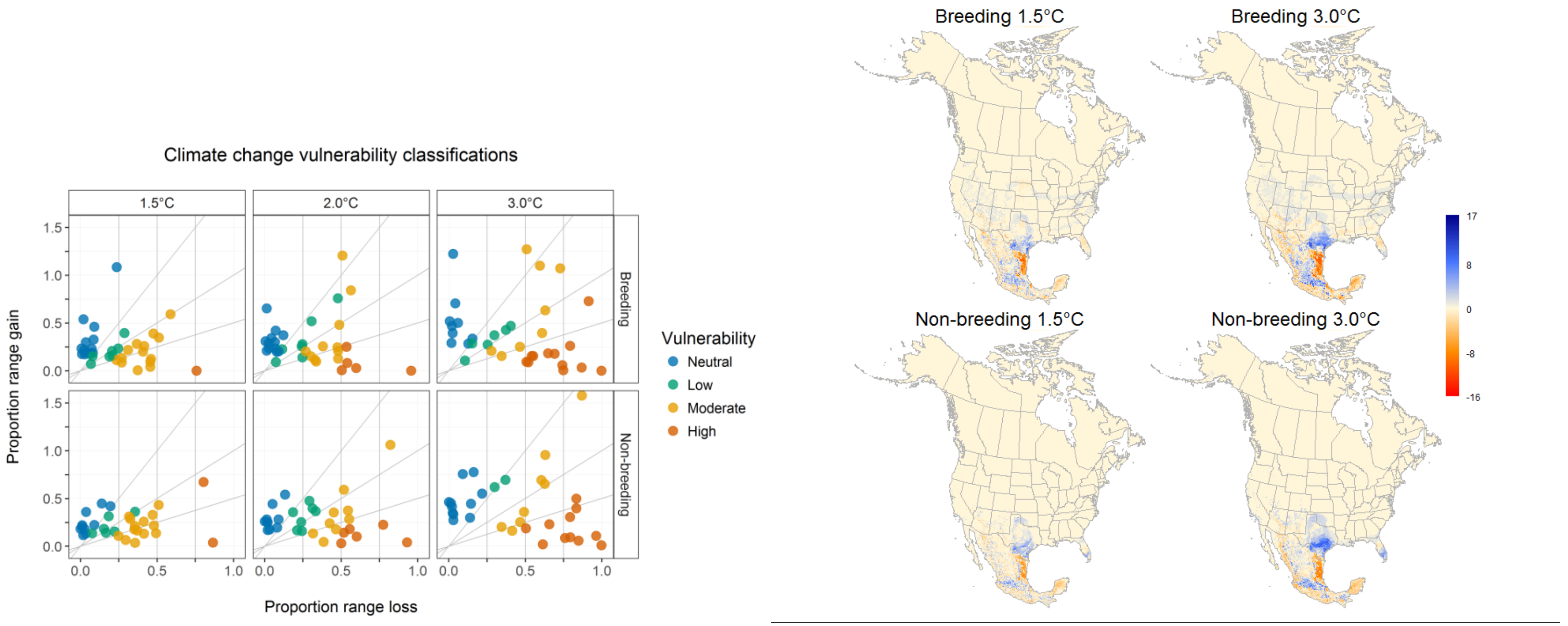


S2.9 Fig. Subtropical Forests bird species vulnerability plot and net change maps.

Subtropical forest bird species results for a) vulnerability of all Subtropical forest species in breeding and non-breeding seasons under 1.5 °C, 2.0 °C and 3.0 °C global warming scenarios, and b) Net gain or loss in number of species at the local community level across Subtropical forest species for the breeding and non-breeding seasons. The scale ranges from net loss (red) to a net gain (blue) of species under 1.5°C warming (left) and 3.0°C warming (right). Net change in the breeding season range from -16 to 17, and from -16 to 17 species in the non-breeding under 3.0°C. Under 1.5°C net change in the breeding seasons ranges from -14 to 15, and from -14 to 13 species in the non-breeding. Net loss is the dominant pattern for subtropical forests species in both the breeding season (median net range loss of -18.1%) and non-breeding season (median range loss of -12.65%), loss in tropical wet and dry forests of Mexico. For this group, we saw species range gains in Texas and the Mexican southern semi-arid highlands and temperate sierras (S3.2 Table). We also saw opposite patterns between seasons in Florida, with loss in the breeding season and gains in the non-breeding season.

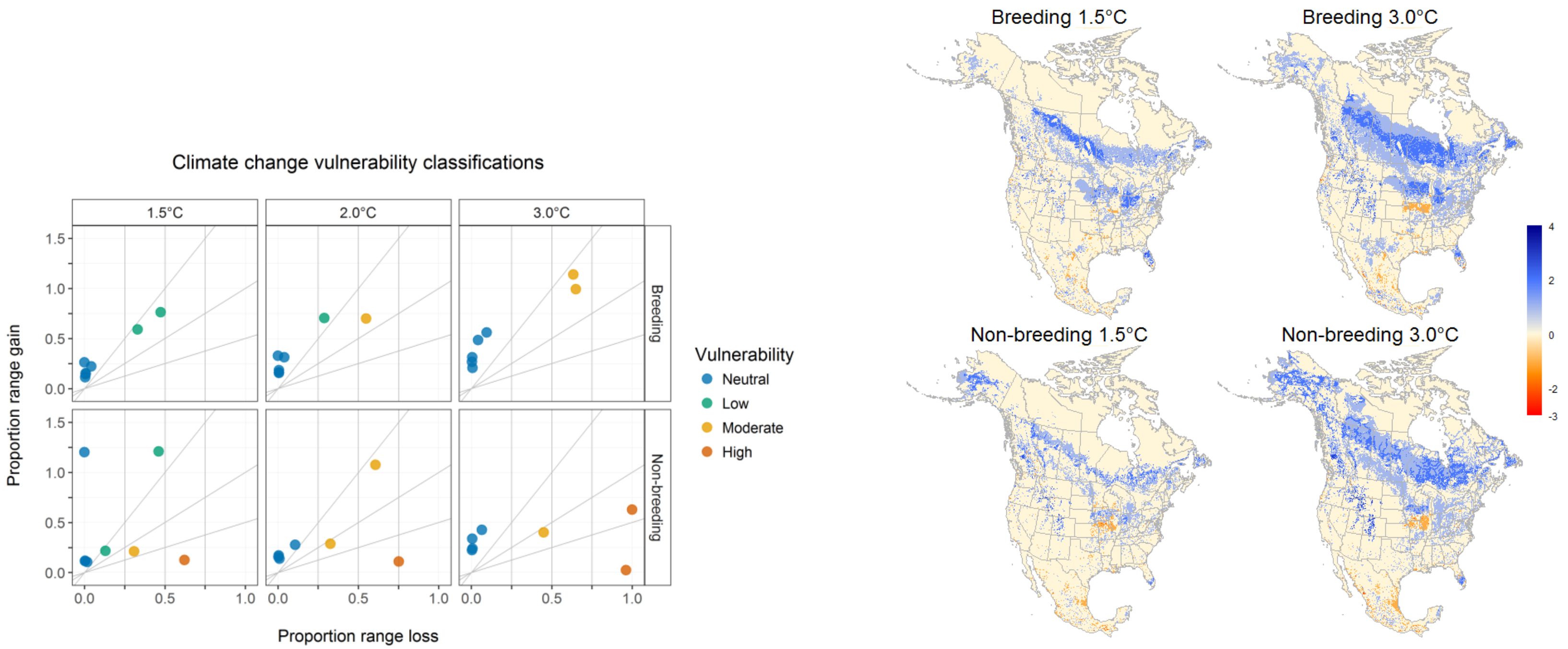


### S2.10 Fig. Urban/Suburban bird species vulnerability plot and net change maps.

Urban/Suburban bird species results for a) vulnerability of all Urban/Suburban species in breeding and non-breeding seasons under 1.5 °C, 2.0 °C and 3.0 °C global warming scenarios, and b) Net gain or loss in number of species at the local community level across Urban/Suburban species for the breeding and non-breeding seasons. The scale ranges from net loss (red) to a net gain (blue) of species under 1.5°C warming (left) and 3.0°C warming (right). Net change in the breeding season range from -3 to 3, and from -3to 4 species in the non-breeding under 3.0°C. Under 1.5°C net change in the breeding seasons ranges from -3 to -3, and from -3 to 4 species in the non-breeding. Urban/Suburban species group have a 45.35% median net range gain in the breeding season (S3.2 Table), although from a small group of eight mostly introduced species. For this group, we see expansions in range gains for most of the Eastern Temperate forest, northern forest and taiga. We also see net gain in the non-breeding season (median net range gain of 32.75) across the same areas.

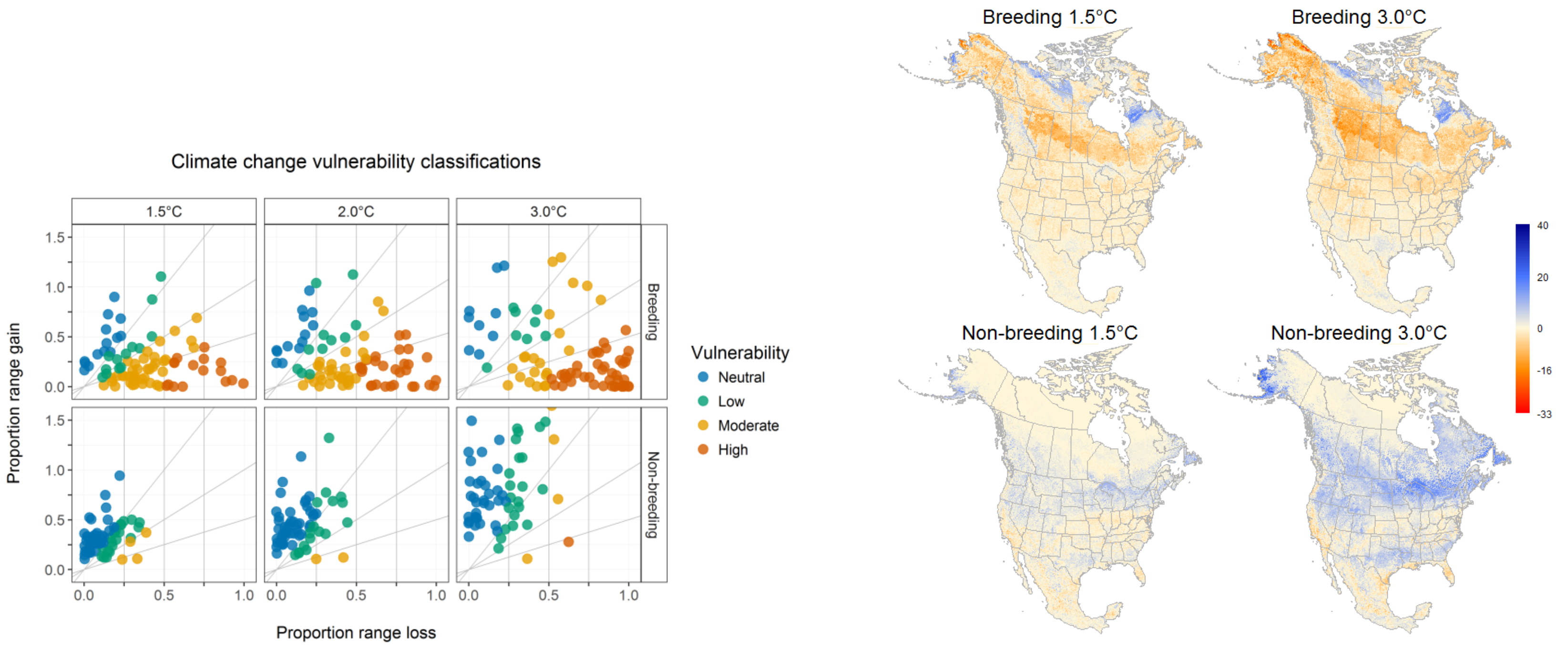


### S2.11 Fig. Waterbirds bird species vulnerability plot and net change maps.

Waterbird species results for a) vulnerability of all Waterbird species in breeding and non-breeding seasons under 1.5 °C, 2.0 °C and 3.0 °C global warming scenarios, and b) Net gain or loss in number of species at the local community level across Waterbird species for the breeding and non-breeding seasons. The scale ranges from net loss (red) to a net gain (blue) of species under 1.5°C warming (left) and 3.0°C warming (right). Net change in the breeding season range from -33 to 20, and from -21 to 40 species in the non-breeding under 3.0°C. Under 1.5°C net change in the breeding seasons ranges from -28 to 23, and from -17 to 24 species in the non-breeding. Waterbird species have a median net loss of -39.5% of their range in the breeding season across much of North America, with strong loss concentrated in the Northern Forest and Taiga (S3.2 Table). This group exhibits strong gains in the non-breeding season (median net gain 53.4%), which are spread across most of North America with the exception of the coastal areas in the southern US. This rate of range gains was the highest out of any group or season (S3.2 Table).

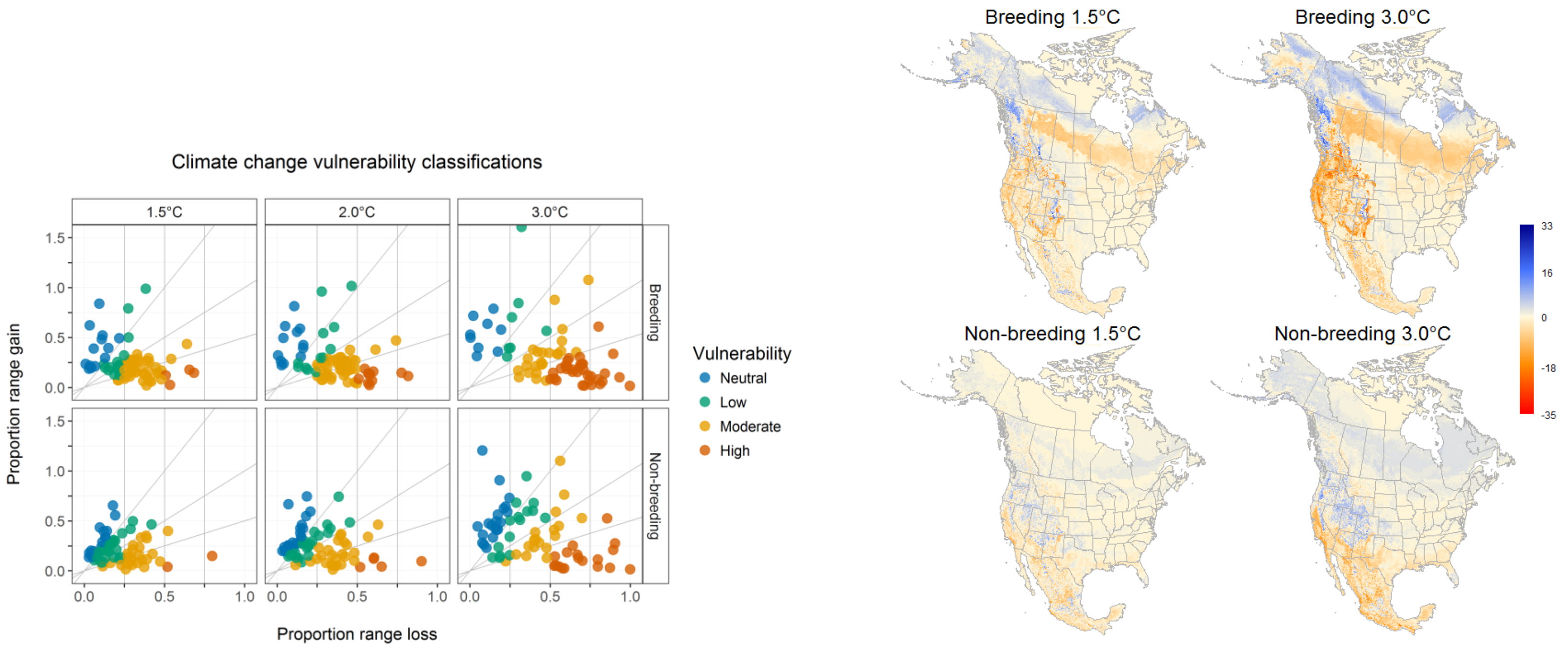


### S2. 12 Fig. Western Forests bird species vulnerability plot and net change maps.

Western forest bird species results for a) vulnerability of all Western forest species in breeding and non-breeding seasons under 1.5 °C, 2.0 °C and 3.0 °C global warming scenarios, and b) Net gain or loss in number of species at the local community level across Western forest species for the breeding and non-breeding seasons. The scale ranges from net loss (red) to a net gain (blue) of species under 1.5°C warming (left) and 3.0°C warming (right). Net change in the breeding season range from -35 to 28, and from -30 to 33 species in the non-breeding under 3.0°C. Under 1.5°C net change in the breeding seasons ranges from -30 to 30, and from -25 to 21 species in the non-breeding. The western forests group is had dominant net loss in both seasons, although less pronounced in the non-breeding season (S3.2 Table). Loss was dominant in Mediterranean California and across Mexico in both seasons, while opposite patterns were seen in the North American deserts, Northern forests, and Northwestern forested Mountains in breeding (loss dominant) and non-breeding (gain dominant).

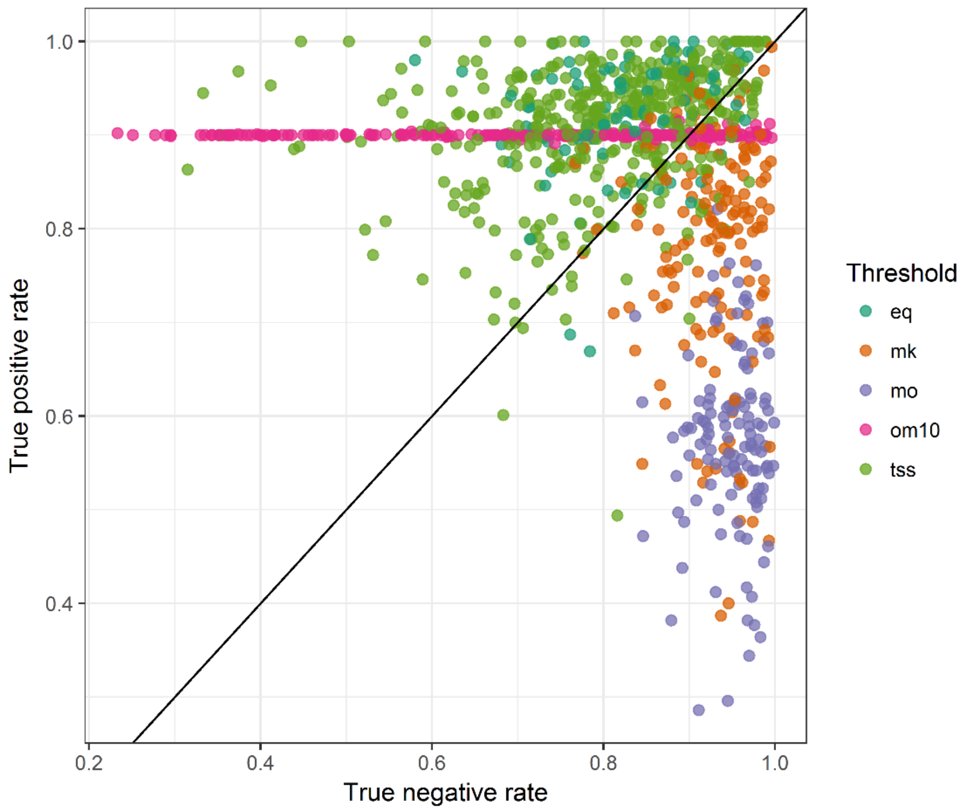


S2.13. Fig. Threshold value for true positive and true negative rates by threshold choice.

Choice of threshold affected current range delineation, with stricter threshold choices (i.e., higher suitability values, e.g. mo, mk) having lower true positive rates and higher true negative rates (i.e. excluded unoccupied and less suitable areas to capture the best possible range, typical of more range-restricted species). In contrast, more relaxed thresholds (i.e., lower suitability values, e.g. min_pred, custom) had higher true positive rates and lower true negative rates (i.e. allowed more of the range to be included, typical of widespread and generalist species). For final choice of threshold based on ecological knowledge and results from the TPR and TNR rates for each species, see S6.

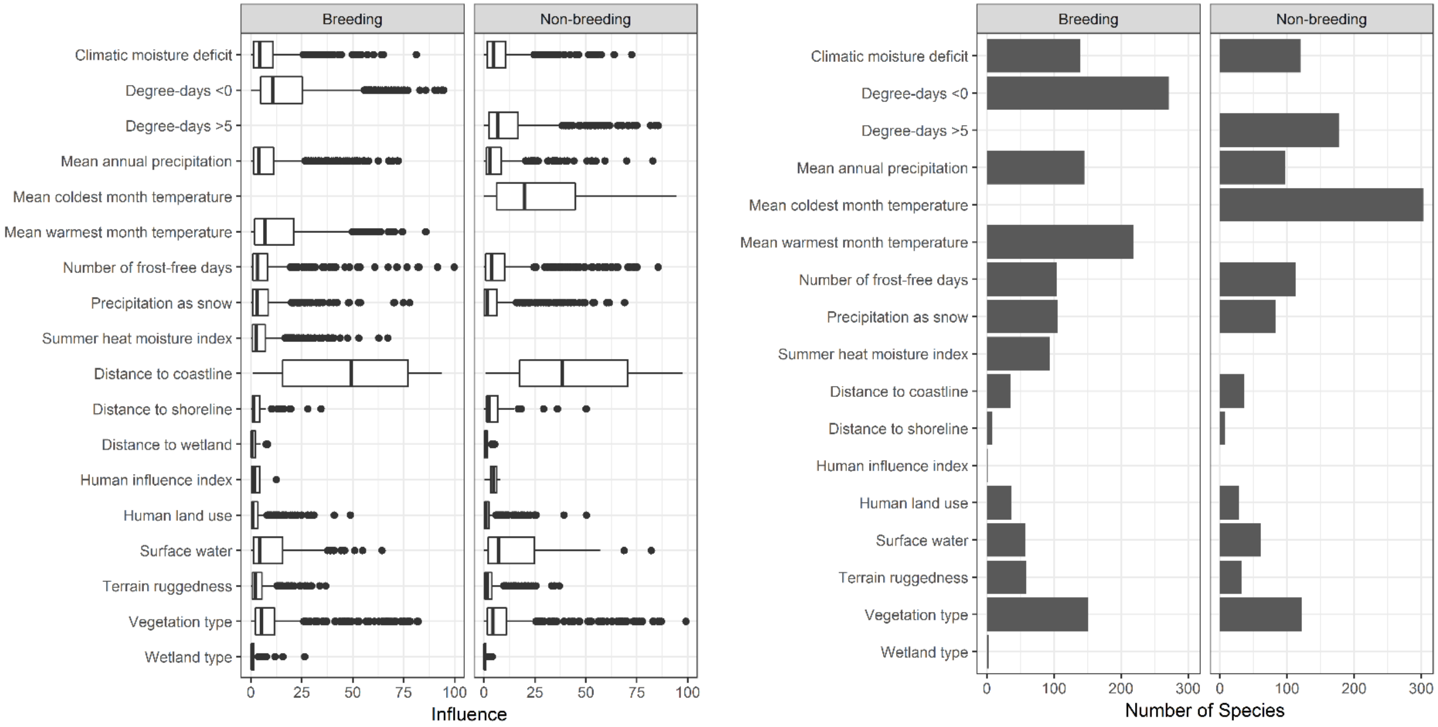


S2.14 Fig. Variable importance (influence) across all species and number of species where each variables was included in the model in the breeding and non-breeding season.
