## Supplementary material for "North American Birds Require Mitigation and Adaptation to Reduce Vulnerability to Climate Change": S3 Supplementary Tables

### S3.1 Table. Species vulnerability by group and scenario

| Breeding Season | | | | | |
|---|---|---|---|---|---|
| **Group** | **Total Species** | **Vulnerable 1.5 (N)** | **Vulnerable 3.0 (N)** | **Vulnerable 1.5 (%)** | **Vulnerable 3.0 (%)** |
| **Arctic** | 16 | 16 | 16 | 100% | 100% |
| **Aridlands** | 69 | 13 | 22 | 19% | 32% |
| **Boreal Forests** | 48 | 36 | 47 | 75% | 98% |
| **Coastal** | 49 | 21 | 24 | 43% | 49% |
| **Eastern Forests** | 69 | 21 | 35 | 30% | 51% |
| **Generalists** | 48 | 5 | 13 | 10% | 27% |
| **Grasslands** | 39 | 15 | 24 | 38% | 62% |
| **Marshlands** | 60 | 11 | 19 | 18% | 32% |
| **Subtropical Forests** | 35 | 15 | 20 | 43% | 57% |
| **Urban/Suburban** | 8 | 0 | 3 | 0% | 38% |
| **Waterbirds** | 83 | 56 | 65 | 67% | 78% |
| **Western Forests** | 73 | 45 | 57 | 62% | 78% |
| **Non-breeding season** | | | | | |
| **Group** | **Total Species** | **Vulnerable 1.5 (N)** | **Vulnerable 3.0 (N)** | **Vulnerable 1.5 (%)** | **Vulnerable 3.0 (%)** |
| **Arctic** | 15 | 5 | 7 | 33% | 47% |
| **Aridland** | 69 | 8 | 19 | 12% | 28% |
| **Boreal Forests** | 38 | 13 | 22 | 34% | 58% |
| **Coastal** | 57 | 10 | 17 | 18% | 30% |
| **Eastern Forests** | 55 | 14 | 17 | 25% | 31% |
| **Generalists** | 45 | 0 | 4 | 0% | 9% |
| **Grasslands** | 35 | 3 | 10 | 9% | 29% |
| **Marshlands** | 57 | 2 | 5 | 4% | 9% |
| **Subtropical Forests** | 32 | 16 | 19 | 50% | 59% |
| **Urban Suburban** | 8 | 2 | 3 | 25% | 38% |
| **Waterbirds** | 63 | 4 | 6 | 6% | 10% |
| **Western Forests** | 71 | 27 | 36 | 38% | 51% |

Number and percent of species in each habitat group considered vulnerable (classified in our analysis as moderate to highly vulnerable) under 1.5 °C and 3.0 °C global warming scenarios, and by season.

**S3.2 Table. Group based median range loss and gain at 1.5°C and 3.0°C by season.**

| Breeding Season | | | | | | |
|---|---|---|---|---|---|---|
| **Group** | **Gain 1.5** | **Loss 1.5** | **Change** | **Gain 3.0** | **Loss 3.0** | **Change** |
| **Arctic** | 10.75 | 39.45 | -28.7 | 4.05 | 78.3 | -74.25 |
| **Aridlands** | 20.8 | 11.3 | 9.5 | 38.7 | 19.2 | 19.5 |
| **Boreal forests** | 25.2 | 36 | -10.8 | 29 | 74.1 | -45.1 |
| **Coastal** | 14.2 | 14.9 | -0.7 | 34.8 | 28.5 | 6.3 |
| **Eastern Forests** | 24.7 | 15.5 | 9.2 | 36.3 | 36.7 | -0.4 |
| **Generalists** | 17.6 | 6.4 | 11.2 | 32.6 | 11.4 | 21.2 |
| **Grasslands** | 23.65 | 24.2 | -0.55 | 37.3 | 51.2 | -13.9 |
| **Marshlands** | 23.8 | 16.85 | 6.95 | 51.6 | 29.65 | 21.95 |
| **Subtropical Forests** | 19.99 | 23.6 | -3.61 | 28.5 | 46.6 | -18.1 |
| **Urban Suburban** | 24.2 | 2.5 | 21.7 | 52.2 | 6.85 | 45.35 |
| **Waterbirds** | 19.7 | 33.5 | -13.8 | 20.7 | 60.2 | -39.5 |
| **Western Forests** | 19.5 | 28.4 | -8.9 | 23.5 | 54 | -30.5 |
| **Non-breeding Season** | | | | | | |
| **Group** | **Gain 1.5** | **Loss 1.5** | **Change** | **Gain 3.0** | **Loss 3.0** | **Change** |
| **Arctic** | 23 | 15.3 | 7.7 | 52.9 | 35.5 | 17.4 |
| **Aridlands** | 24.8 | 7.9 | 16.9 | 46 | 13.4 | 32.6 |
| **Boreal forests** | 19.7 | 17.3 | 2.4 | 40.7 | 42.3 | -1.6 |
| **Coastal** | 26.1 | 9.9 | 16.2 | 60.2 | 25 | 35.2 |
| **Eastern Forests** | 18.3 | 10.4 | 7.9 | 44.3 | 22.3 | 22 |
| **Generalists** | 19.1 | 7.7 | 11.4 | 47.7 | 11.4 | 36.3 |
| **Grasslands** | 23.5 | 11.1 | 12.4 | 47.1 | 23.1 | 24 |
| **Marshlands** | 24.7 | 6.7 | 18 | 61.1 | 13.4 | 47.7 |
| **Subtropical Forests** | 19.2 | 27.25 | -8.05 | 35.3 | 47.95 | -12.65 |
| **Urban Suburban** | 16.75 | 7.4 | 9.35 | 36.9 | 4.15 | 32.75 |
| **Waterbirds** | 29.9 | 11.6 | 18.3 | 72.1 | 18.7 | 53.4 |
| **Western Forests** | 17.7 | 18.4 | -0.7 | 33.2 | 36.7 | -3.5 |

**S3.3 Table. Variable importance by bird habitat group and season**

| Breeding Season | | | | | | | | | | |
|---|---|---|---|---|---|---|---|---|---|---|
| | Variable | | | | | | | | | |
| Group | Anthropogenic land use | Degree-days <0 | Climatic moisture deficit | Mean annual precipitation | Mean warmest month temperature | Number of frost-free days | Precipitation as snow | Summer heat moisture index | Topographic roughness | Vegetation type |
| Arctic | 0.58 | 14.95 | 8.73 | 2.02 | 39.81 | 5.89 | 2.28 | 1.21 | 2.70 | 3.42 |
| Aridlands | 0.55 | 14.85 | 7.85 | 5.42 | 1.65 | 1.41 | 2.74 | 6.45 | 1.09 | 4.00 |
| Boreal forests | 2.96 | 13.99 | 12.32 | 1.41 | 31.17 | 3.72 | 4.42 | 1.52 | 0.94 | 7.28 |
| Coastal | 0.43 | 3.30 | 0.65 | 0.70 | 2.17 | 2.15 | 0.50 | 0.44 | 0.53 | 0.80 |
| Eastern Forests | 8.29 | 8.38 | 3.49 | 11.71 | 17.54 | 4.51 | 8.05 | 5.02 | 3.66 | 4.22 |
| Generalists | 4.86 | 12.38 | 8.20 | 6.17 | 6.96 | 7.45 | 7.85 | 4.84 | 3.13 | 4.88 |
| Grasslands | 0.90 | 11.88 | 5.01 | 17.98 | 6.29 | 4.93 | 6.04 | 3.19 | 4.10 | 3.78 |
| Marshlands | 0.59 | 12.43 | 1.37 | 8.41 | 3.28 | 3.47 | 2.38 | 1.24 | 6.00 | 4.84 |
| Subtropical Forests | 0.45 | 20.20 | 1.91 | 1.09 | 1.38 | 4.14 | 0.20 | 3.16 | 0.69 | 7.37 |
| Urban Suburban | 1.45 | 12.97 | 3.06 | 2.09 | 2.52 | 2.30 | 0.08 | 2.24 | 0.32 | 4.84 |
| Waterbirds | 1.15 | 8.11 | 4.86 | 2.97 | 14.80 | 3.00 | 1.95 | 2.09 | 2.26 | 6.19 |
| Western Forests | 0.89 | 20.81 | 3.66 | 3.09 | 6.16 | 3.63 | 3.39 | 3.00 | 3.15 | 7.08 |
| Non-breeding season | | | | | | | | | | |
| | Variable | | | | | | | | | |
| Group | Anthropogenic land use | Degree-days >5 | Climatic moisture deficit | Mean annual precipitation | Mean coldest month temperature | Number of frost-free days | Precipitation as snow | | Topographic roughness | Vegetation type |
| Arctic | 1.71 | 13.10 | 2.92 | 3.26 | 30.96 | 5.23 | 2.03 | | 2.81 | 3.91 |

| | | | | | | | | | | |
|---|---|---|---|---|---|---|---|---|---|---|
| **Aridlands** | 0.88 | 5.13 | 8.05 | 9.37 | 16.76 | 2.53 | 3.62 | | 0.94 | 8.57 |
| **Boreal forests** | 2.02 | 33.21 | 4.16 | 1.96 | 14.70 | 2.52 | 2.10 | | 0.52 | 4.54 |
| **Coastal** | 0.47 | 6.73 | 1.53 | 0.60 | 6.50 | 3.84 | 0.34 | | 0.29 | 0.52 |
| **Eastern Forests** | 1.54 | 10.10 | 3.59 | 6.41 | 30.62 | 3.63 | 1.39 | | 2.07 | 4.33 |
| **Generalists** | 3.40 | 14.26 | 5.80 | 4.94 | 22.27 | 5.94 | 5.17 | | 2.23 | 5.06 |
| **Grasslands** | 2.99 | 7.41 | 8.54 | 6.09 | 6.00 | 6.66 | 5.78 | | 4.08 | 4.98 |
| **Marshlands** | 0.94 | 5.07 | 3.01 | 1.35 | 21.83 | 5.11 | 1.53 | | 2.04 | 2.78 |
| **Subtropical Forests** | 0.42 | 4.42 | 2.28 | 2.72 | 62.49 | 1.29 | 1.05 | | 0.73 | 4.01 |
| **Urban Suburban** | 2.39 | 7.91 | 3.89 | 8.10 | 12.06 | 2.41 | 2.99 | | 4.04 | 2.28 |
| **Waterbirds** | 0.83 | 7.45 | 4.52 | 1.75 | 14.21 | 7.77 | 1.92 | | 0.93 | 2.11 |
| **Western Forests** | 0.62 | 6.41 | 7.67 | 2.94 | 28.54 | 2.38 | 0.83 | | 3.10 | 5.92 |

**S3.4 Table. Habitat Bird Group specific variable importance by group and season**

| **Breeding Season** | | | | | | |
|---|---|---|---|---|---|---|
| | **Variable** | | | | | |
| **Group** | **Wetland Type** | **Distance to Wetland** | **Distance to coastline** | **Distance to shoreline** | **Surface Water occurrence** | **Human Influence Index** |
| **Coastal** | | | 59.58 | | 1.57 | |
| **Marshlands** | 0.81 | | | 2.20 | 9.96 | |
| **Urban Suburban** | | | | | | 5.03 |
| **Waterbirds** | 0.58 | 0.61 | | | 3.78 | |
| **Non-breeding season** | | | | | | |
| | **Variable** | | | | | |
| **Group** | **Wetland Type** | **Distance to Wetland** | **Distance to coastline** | **Distance to shoreline** | **Surface Water occurrence** | **Human Influence Index** |
| **Coastal** | | | 41.38 | | 1.94 | |
| **Marshlands** | 0.66 | | | 2.43 | 7.12 | |
| **Urban Suburban** | | | | | | 6.18 |
| **Waterbirds** | 0.17 | 0.72 | | | 19.81 | |
